# Protrusion force and cell-cell adhesion-induced polarity alignment govern collective migration modes

**DOI:** 10.1101/2024.09.18.613706

**Authors:** Huijing Wang, Catalina Ardila, Ajita Jindal, Vaishali Aggarwal, Weikang Wang, Jonathan Vande Geest, Yi Jiang, Jianhua Xing, Shilpa Sant

## Abstract

Collective migration refers to the coordinated movement of cells as a single unit during migration. While collective migration enhances invasive and metastatic potential in cancer, the mechanisms driving this behavior and regulating tumor migration plasticity remain poorly understood. This study provides a mechanistic model explaining the emergence of different modes of collective migration under hypoxia-induced secretome. We focus on the interplay between cellular protrusion force and cell-cell adhesion using collectively migrating three-dimensional microtumors as models with well-defined microenvironments. Large microtumors show directional migration due to intrinsic hypoxia, while small microtumors exhibit radial migration when exposed to hypoxic secretome. Here, we developed an in silico **m**ulti-**s**cale **m**icrotumor **m**odel (MSMM) based on the cellular Potts model and implemented in CompuCell3D to elucidate underlying mechanisms. We identified distinct migration modes within specific regions of protrusion force and cell-cell adhesion parameter space and studied these modes using in vitro experimental microtumor models. We show that sufficient cellular protrusion force is crucial for radial and directional collective microtumor migration. Radial migration emerges when sufficient cellular protrusion force is generated, driving neighboring cells to move collectively in diverse directions. Within migrating tumors, strong cell-cell adhesion enhances the alignment of cell polarity, breaking the symmetric angular distribution of protrusion forces and leading to directional microtumor migration. The integrated results from the experimental and computational models provide fundamental insights into collective migration in response to different microenvironmental stimuli. Our computational and experimental models can adapt to various scenarios, providing valuable insights into cancer migration mechanisms.

**Statement of Significance:** While single-cell metastasis is well-studied, mechanisms of collective cluster migration are less understood. Significant challenges include the lack of a fundamental physics perspective on collective cluster migration mechanisms and suitable physiologically relevant three-dimensional (3D) in vitro models that can recapitulate collective cluster migration. In this article, we developed a computational model depicting microtumor migration behaviors. Collective cell migration, with varying correlation lengths, exhibits different migratory modes such as directional and radial migration. These modes are predicted by in silico models and confirmed using experimental microtumor models. Machine learning methods were exploited to identify migratory modes. Our computational and experimental models are flexible in various circumstances, offering insights into cancer migration mechanisms.

## Introduction

Cancer metastasis is a multi-step process that involves migration, invasion, and dissemination of tumor cells from the primary tumor site to the distant organs to establish secondary metastases (1). There is growing evidence that cancer metastases and invasion are driven by the movement of small tumor cell clusters rather than individual cells (2). This phenomenon is called collective migration, which refers to the coordinated movement of cells as a single unit due to strong cell-cell adhesion forces maintained by cell-cell junctions (3–6). The role of collective cell migration in cancer metastasis was first reported in the 1950s in a study that showed the presence of both individual cells and collective tumor clusters in a cancer patient’s blood. The researchers established that these circulating tumor cell (CTC) clusters originate from solid tumors and are shed into the bloodstream (7,8). It is also reported that the tumor clusters move more efficiently through the bloodstream when disseminated as a group of cells (2) and can retain their epithelial cell adhesion machinery that contributes significantly to metastatic progression (9). Although collective migration is known to drive several other biological processes, such as wound healing and embryogenesis, the underlying mechanisms behind various modes of collective migration are poorly explored.

Symmetry breaking is an essential process underlying cellular fate decisions, such as polarization, migration, and division. For example, a local asymmetric distribution of Ras-GTP signal triggers cytoskeletal rearrangement, thus establishing single cell polarity in eukaryotic amoeboid cells for migration (10–12). However, the mechanistic interactions for spontaneous symmetry breaking in the collective migration of tumor clusters have not been fully elucidated (11,13). Understanding such tumor migration mechanisms, including symmetry breaking, could help design new cancer treatments; for example, regulators or key components that control these mechanisms could be targeted.

A major hurdle in studying collective migration in tumor clusters is the lack of suitable physiologically relevant three-dimensional (3D) *in vitro* models that can recapitulate collective cluster migration. *In vitro* 2D assays such as Boyden chamber assays or the scratch assay (14,15) have been widely used to study single cell migration and invasion; however, this technique is not suitable for studying collective cluster migration (16) because these 2D assays cannot quantify collective tumor migration, the tumor shape, or cell-cell interactions (17). To unravel the influence of a hypoxic tumor microenvironment on the emergence of migratory phenotype, we have previously developed a hydrogel device with microwell arrays to generate uniform-sized large (600 µm) and small (150 µm) microtumors from the same non-invasive parental T47D breast cancer cells (18–25). We have shown that the large microtumors develop hypoxic signatures as early as day 1 and exhibit collective migration (19), where whole tumor clusters migrate out of the hydrogel microwell. On the other hand, the non-hypoxic small microtumors do not migrate over the culture period. However, when treated with the secretome from large hypoxic microtumors, the non-migratory small microtumors exhibit starburst-like radial migration (21). Thus, microtumors made of the same parental T47D cells showed plasticity in collective migration, where the directional and radial modes of collective migration emerged in response to two different microenvironmental stimuli, tumor-intrinsic hypoxia and hypoxic secretome, respectively. In this work, we set out to investigate plausible mechanisms underlying the emergence of these distinct migratory phenotypes by using pharmacological perturbations and mathematical approaches.

Tumor migration results from a series of biological processes (26). It is often associated with mechanical cues in the microenvironment due to shear stresses, cell-cell and cell-ECM interactions (27), where tumor cells respond to signaling pathways such as Rho GTPases by polarizing and extending actin polymerization-driven cell membrane protrusions towards the extracellular cues, such as growth factors or chemokines (28–30). Tumor cell plasticity or increased metabolism, among others, have been proposed as potential mechanisms for tumor migration, including single and collective cell migration (31). However, there is no consensus on how different migratory phenotypes emerge. In this work, we investigated the interplay between protrusion force and cell-cell adhesion to explain the plasticity of collective tumor migration in response to different microenvironmental stimuli. To elucidate potential mechanisms underlying the emergence of different tumor migration modes, such as directional cluster migration and radial migration, we designed the in silico **m**ulti-**s**cale **m**icrotumor **m**odel (MSMM). The MSMM is a cellular Potts model and implemented in CompuCell3D (32,33). It is informed by *in vitro* 3D microtumor experiments, and incorporates cellular mechanics, such as cell shape distortion, force-driven motility, and cell-cell adhesion. By fine-tuning the protrusion and cell-cell adhesion forces in our MSMM, we can recapitulate both, the directional and the radial migration modes observed experimentally in response to different stimuli. Combining pharmacological perturbations and mathematical approaches, we demonstrate that threshold protrusion force is necessary for collective migration. In contrast, distinct migration modes emerge due to competition between protrusion and cell-cell adhesion forces.

## Materials and Methods

### Cell lines and cell culture

The T47D breast cancer cell line (RRID:CVCL_0553) was purchased from the American Type Culture Collection (ATCC, Manassas, VA, USA). Media for cell culture was obtained from Corning® unless specified. The cells were passaged and maintained in T75 flasks using Dulbecco’s modified Eagle’s medium (DMEM) (MT10013CV, Corning®, Manassas, VA, USA) supplemented with 10% fetal bovine serum (FBS) (S11250, Atlanta Biologicals, Flowery Branch, GA, USA), and 1% penicillin-streptomycin (300-002-CI, Corning®, Manassas, VA, USA) in a humidified atmosphere at 37°C and 5% CO2. Cells were maintained between 60-70% confluency for further passaging or seeding into hydrogel microwell devices. Cell line authentication was done by the University of Arizona, Genetics Core (UAGC, Chandler, AZ) using PowerPlex16HS PCR Kit as previously described (21).

### Microtumor fabrication

Uniform-sized microtumors of 150 µm diameter were fabricated by seeding T47D cells in Polyethylene glycol dimethacrylate (PEGDMA) hydrogel microwell devices as previously described (18,19,21,22). Briefly, polydimethylsiloxane (PDMS) stamps (1×1 cm^2^) containing posts of 150 µm in diameter and 150 µm in height were used to produce microwell devices. A solution of polyethylene glycol dimethacrylate (20% w/v, PEGDMA, 1000Da, Polysciences Inc, Warrington, PA, USA) containing photoinitiator (Irgacure-1959, 1% w/v, Ciba AG CH-4002, Basel, Switzerland) was crosslinked under the PDMS stamps using OmniCure S2000 curing station (200W Lamp, 5W/cm^2^, EXFO, Mississauga, Canada). The fabricated microwell hydrogel devices were sterilized by submerging them in 70% ethanol under UV light for 1h. Sterilized hydrogel devices were washed thrice with Dulbecco’s phosphate buffered saline (DPBS) without calcium and magnesium (Corning™, Manassas, VA, USA, catalog #21-031-CV). For cell seeding, a suspension of 1 x10^6^ T47D cells (RRID:CVCL_0553) in 50 µL of growth media was dropped on each hydrogel microwell device. Cells were allowed to settle in the microwells for 15 min. The cells that did not settle into the microwells were removed by gentle washing with DPBS. The cell-seeded devices were cultured at 37 °C and 5% CO2 in a humidified incubator. On day 1 and day 2, 50% of the culture media was replaced with an equal volume of fresh media. To study migration induced by tumor-secreted factors, 150µm microtumors were treated with conditioned media (CM) from hypoxia-rich larger (600µm) microtumors starting from day 3 until day 6 (referred to as **150CM** hereafter), with the replacement of 50% media with an equal amount of CM every day as described in our previous studies (21). Control 150µm microtumors (referred to as **150** hereafter) were cultured similarly for 6 days with a replacement of 50% media with fresh media every day, and these microtumors served as non-migratory controls.

### Assessment of microtumor migration

The aggressiveness of the migratory phenotype of microtumors was assessed by quantifying the percentage of migrating tumors from each microwell hydrogel device (n=3) and the migration distance. The distance of migration was estimated in ImageJ (RRID:SCR_003070) by measuring the average length of at least 10 straight lines drawn perpendicular to the boundary of the well up to the leading edge of the microtumor, covering the entire migratory front of at least 10 migrating microtumors per group.

### Microarrays and bioinformatics analysis

Microarray data was collected as described previously (23). Briefly, at the end of six days in culture, RNA was isolated from 150D6 (control non-migratory microtumors) and 150CM microtumors using GenElute™ Mammalian Total RNA Miniprep Kit (Catalog No. RTN70-1KT, Sigma-Aldrich, St. Louis, MO, USA) and DNA cleanup was done by On-Column DNase I Digestion Set (Catalog No. DNASE70-1SET, Sigma-Aldrich, St. Louis, MO, USA) as per manufacturer’s protocols. RNA quantity was measured by absorbance ratio at 260/280 nm, and integrity was verified on 1% agarose gel. cDNA preparation, hybridization to GeneChips, scanning, and first-level data analysis were performed at the University of Pittsburgh HSCRF Genomics Research Core as described previously (23). Microarray data is deposited in the Gene Expression Omnibus database at the National Center for Biotechnology Information (GSE166211, RRID:SCR_005012). Differential gene expression in migratory 150CM induced by the secretome was analyzed by normalization with the 150D6 (150CM *versus* 150D6). Transcriptome Analysis Console (TAC 4.0, ThermoFisher Scientific, Waltham, MA, USA) was used to perform statistical analysis on the comparison using one-way ANOVA. The initial assessment of the microarray expression data was performed by hierarchical clustering using a cut-off *p*-value of 0.01. All the genes with False Discovery Rate (FDR) ≤ 0.05, *p-value* ≤ 0.05, and fold change ± 2 were considered significantly different in gene expression and were selected for further analysis. Gene Set Enrichment Analysis (GSEA) was carried out for Gene Ontology (GO) enrichment of Biological Processes (BP) using the GSEA software with minimum size cut-off =5 (UC San Diego and Broad Institute, San Diego, CA, USA) as discussed previously (23). We performed the analysis of differentially enriched GO: BP related to ‘*actin polymerization*’, ‘*cell-cell adhesion*’, ‘*cell junction*’, ‘*adherence junction*’, ‘*cell polarity*’, and ‘*cytoskeleton reorganization*’ to capture phenotypic differences in directionally migrating 600D6 and radially migrating 150CM microtumors compared to non-migratory 150 µm microtumors.

### Microtumor inhibition studies

Based on our microarray data analysis, we used inhibitors of myosin and AKT2 molecules to investigate their role in developing collective migration phenotypes. All the inhibition treatments started on Day 3, along with the first CM supplement. Myosin light chain (MLC) inhibition was achieved by treating 150CM microtumors with 50 µM of Blebbistatin (Catalog No. 50-101-3165, Apexbio Technology LLC, Houston, TX, USA). Blebbistatin is a cell-permeable, selective pharmacological inhibitor of myosin II (34,35). Complete solubilization of Blebbistatin in either growth media or CM was achieved by 2h sonication in an ultrasonic bath at 37°C and 40kHz. AKT Serine/Threonine Kinase 2 (AKT2) was inhibited chemically by the treatment of 150CM microtumors with growth media CM containing CCT128930 (Catalog No. 50-190-5165, Apexbio Technology LLC, Houston, TX, USA) at a final concentration of 25 µM. CCT128930 is an ATP-competitive selective pharmacological inhibitor of AKT2 and its downstream targets (36,37). Untreated 150CM microtumors were used as controls.

### Immunostaining and confocal microscopy

Microtumors were fixed with 4% paraformaldehyde, permeabilized with 0.1% Triton X-100 in 1X PBS, and blocked with 3% BSA in 1X DPBS containing 0.1% Triton X-100. Microtumors were stained for myosin light chain (MLC) (#109061-1-AP, Proteintech, Rosemont, IL, USA), AKT (#4691, Cell Signaling Technology, Danvers, Massachusetts, USA), phospho-AKT (pAKT) (#4060; Cell Signaling Technology, Danvers, Massachusetts, USA) by incubating the samples with respective primary antibodies overnight at 4°C. Samples were then washed with 1X DPBS and incubated with secondary antibody Alexa Fluor 488 goat anti-rabbit (#A11008, Invitrogen Carlsbad, California, USA) for 2h. Samples were rewashed with 1X DPBS and incubated with a Hoechst solution in 1X DPBS for 24h. The stained samples were washed with 1X DPBS and mounted in glycerol on microscopy slides (#6776214, Thermo Scientific, Waltham, MA, USA). Confocal imaging was carried out on the Olympus FV1000 inverted confocal microscope at the University of Pittsburgh, Department of Pharmaceutical Sciences, using a step size of 5 μm. Maximum intensity Z-projections and intensity plot profiles were generated using FIJI software (ImageJ) (RRID:SCR_002285 and RRID:SCR_003070).

### Statistical Analysis

All the values are presented as mean ± standard deviation. A paired t-test was used to compare inhibitor-treated and untreated control groups statistically. Two-way ANOVA was used to compare the level of significance in the extent of microtumor migration between the inhibitor-treated *versus* untreated control group. Sample sizes for each study were indicated in the respective figure legends. A *p-*value less than 0.05 was considered significant.

### Multi-scale microtumor model energy terms

In the cellular Potts model, to mimic the extension and retraction of the cell, the algorithm randomly selects a lattice site and attempts to copy its index to a random neighboring lattice. The MSMM represents cells on a 256 × 256 lattice, where cells are defined as a set of pixels, and different cells do not overlap. Each cell is constrained by surface energy, boundary energy, and energy from the protrusion force (**Supplementary Table S1**). Cell spatial dimensions were defined based on the assumption that the cells are spherical, with a radius of 4 pixels. This default assumption is based on the fact that breast cancer cell T47D is spherical before it attaches to a surface or after finishing a cell cycle. MSMM does not have a defined “well” comparing the experimental setup. In the simulation, 138 cells were initialized uniformly randomly in a center disc with radius of 33. This number closely matches the initial cell count seeded in a 150 µm microwell. After relaxation, the cell blob has a diameter of roughly 100 pixels, equating to 1.5 µm/pixel.

A global effective energy *H* consists of cell constraints, such as area and perimeter, cell-cell interactions, cell-medium interactions, and protrusion force energy. The global effective energy *H* (38) function is defined as:

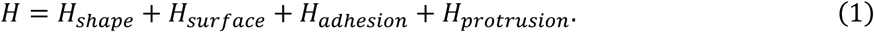

For a cell with a unique index of *σ*, term *H_shape_* comes from cell shape constraints and is defined as:

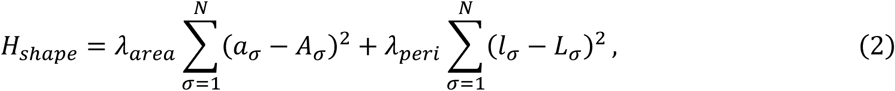

where *λ*_*area*_ and *λ*_*peri*_ are parameters that describe cell elasticity. The *σ* is the cell identity. The total number of cells in the simulation is *N*. *a*_*σ*_ and *l*_*σ*_ are the area and perimeter for cell *σ*, and *A*_*σ*_ and *L*_*σ*_ are the target area and perimeter corresponding to *a*_*σ*_ and *l*_*σ*_.

The second term, *H_surfaca_* in Eq. 1, describes the surface properties for cell-cell, cell-medium, and medium-medium contacts as:

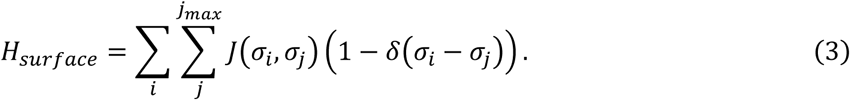

The outer sum of the surface energy term *i* iterates over all cell-lattice pixels. The model refers to the contacted pixels as 1^st^ order pixels and pixels further away as 2^nd^, 3^rd^ order, and so on. The inner sum (over *j*) iterates over all pixels in the neighborhood of *i* up to order *j*_*max*_. *J*(*σ*_*i*_, *σ*_*j*_) is the interface energy per unit contact area defined in the model. The term *δ*(*σ*_*i*_ − *σ*_*j*_) is Kronecker delta that only allows interactions between pairs of lattices belonging to different cells.

The third term on the right-hand side of Eq. 1, *H*_*adhesion*_, describes the adhesion energy, defined as:

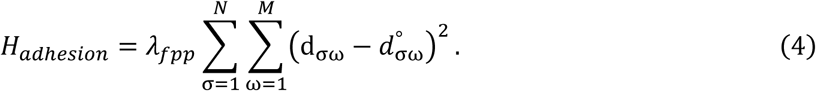

The *H*_*adhesion*_ represents adherens junctions (AJ) that link cells; disrupting the AJs results in loss of cell-cell contacts. In cancer cells, catenins in the cytoplasm bind to cytoskeletal components, such as actin filaments and microtubules to form AJs (39). This adhesion linkage is described as strings that link cell centers using the FocalPointPlasticity (FPP) plugin in CompuCell3D (RRID:SCR_003052). In CompuCell3D, a neighboring cell is defined as an adjacent cell that has a common surface area with the cell in mind. For cell *σ*, *M* represents the count of neighboring cells surrounding *σ*. The d_σω_ is the distance between the centers of mass of cells *σ* and ω, and 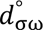 is a target length corresponding to d_σω_. The *λ*_*fpp*_ is a constraint parameter that describes the strength of the focal point links.

The last term, *H_protrusion_* in Eq. 1 describes the energy term simulating protrusion force:

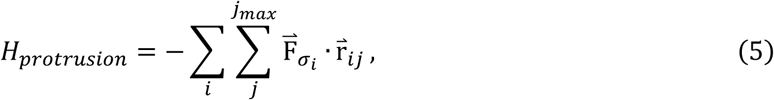

Again, in the outer sum, *i* iterates over all cell-lattice pixels. The inner sum (over *j*) iterates over all pixels in the neighborhood of *i* up to order *j_max_*. *F⃑*_*σi*_ is the protrusion driving force for attempt copy from pixel i to j for cell *σ*, and *r⃑*_*ij*_ is the distance vector (40). *F⃑*_*σi*_ is regulated by single cell protrusion force regulator *R*, and its mathematical expression is given later (Eq. 10).

Conventionally, the multi-scale microtumor model (MSMM) accepts a Monte Carlo move for lattice *u* copy to *v* with the Boltzmann probability *p*(*u* → *v*) as shown in Eq. 6.

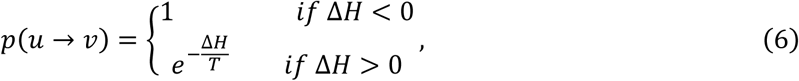

Where, Δ*H* = *H*_*v*_ − *H*_*u*_ is the attempted move contribution to total effective energy. The effective temperature *T*(> 0) is a parameter to control the move acceptance probability, with the acceptance probability *p*(*u* → *v*) increases as *T* increases. We use a constant *T* = 25 for all simulations.

### Secretome signals diffuse into tumor cells

Secretome induces radial migration for 150µm microtumors (**Figure 1b**) by activating an unidentified protein regulator R. There are many potential candidate proteins *R* that can regulate cell cytoskeleton remodeling and generating protrusion force, such as Rac1-GTP, a Rho GTPases family protein typically residing at the migration front, which governs the formation of lamellipodia, or myosin II, a motor protein that binds F-actin to generate contraction force, and so on. The secretome uptake is simulated as a diffusion field for 150µm microtumors treated with the conditioned media (150CM) from hypoxic tumors. The medium serves as the boundary for the secretome signal field, where every pixel has a constant value of 1. The secretome signal diffuses into the tumor cells (both protrusive cells and non-protrusive cells), following the function for location x⃑ :

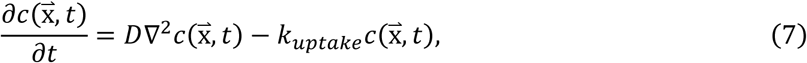

**Figure 1.**
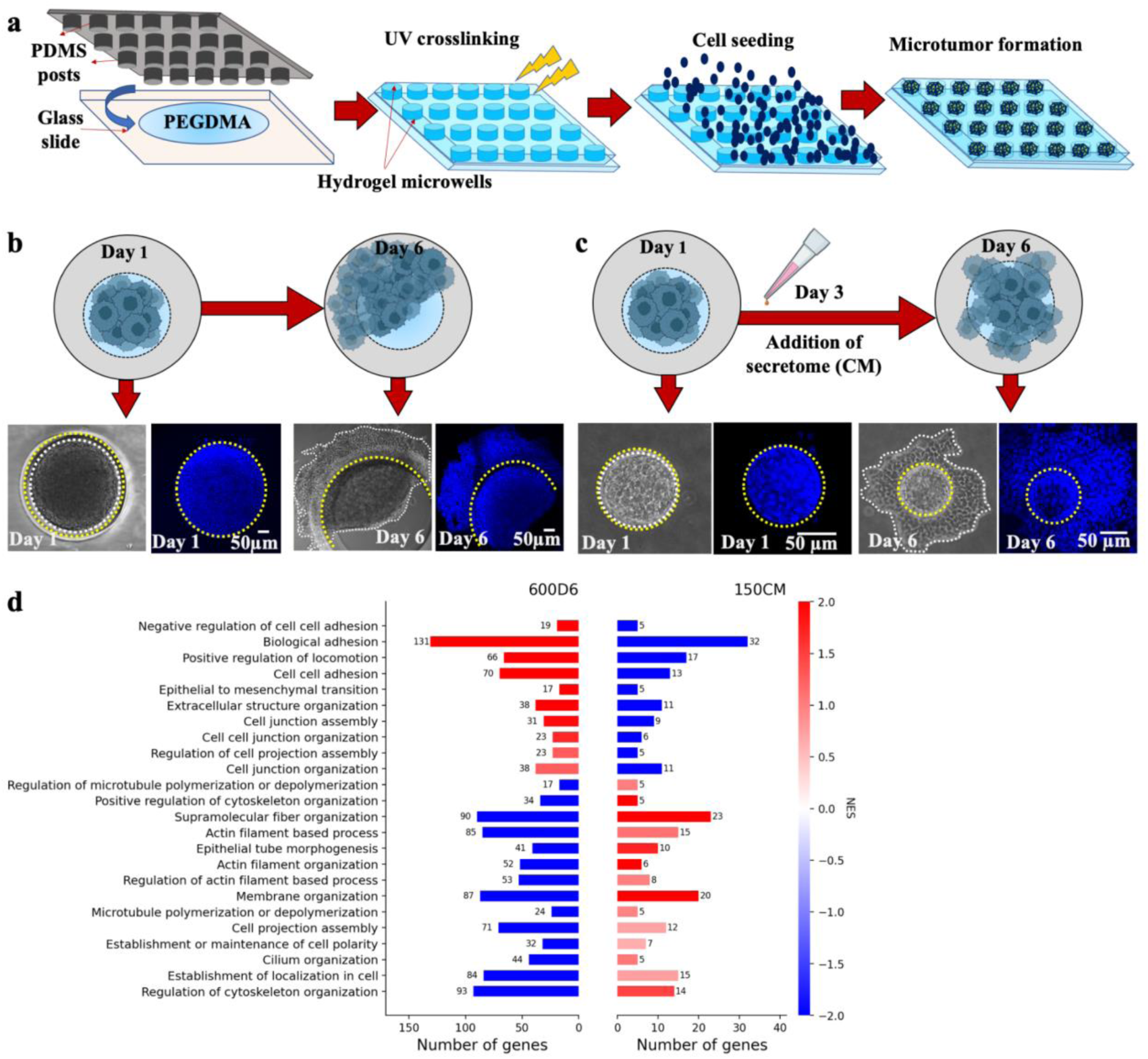
Tumor-intrinsic hypoxia induces directional migration in large microtumors, while hypoxic secretome induces radial migration in small microtumors. a. Schematic representation of microfabrication of uniform-sized T47D microtumors on hydrogel microarrays. b. Hypoxic 600 μm microtumors exhibit directional migration on day 6, scale bar: 50 μm c. Treatment of non-migratory 150 μm tumors with hypoxic secretome (CM) gives rise to radial starburst-like migratory phenotype. Scale bar: 50 μm d. Paired gene set enrichment analysis (GSEA) for GO terms of biological processes related to the biophysical mechanism of microtumor migration, where the left bar plot shows enriched BP for directional migration (600D6) and the right bar plot shows enriched BP for radial migration (150CM). The analysis was done on differentially expressed genes using non-migratory 150D6 as a control group. The normalized enrichment scores (NES) for biological processes are color-coded. The names of biological processes are labeled on the y-axis. The x-axis shows the number of differentially expressed genes for each GO term.

where *c*(x⃑, *t*) is the concentration of secretome signal at position x⃑ and time *t*. *D* is the diffusion coefficient, and *k*_*uptake*_ is the pixel-wise uptake rate by cell at position x⃑ and time *t*.

Thus, for cell *σ* at the time *t*, the production rate *s* of protein *R*, which is dependent on a single cell secretome signal, can be obtained by

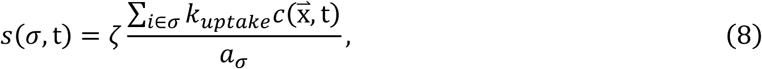

where *i* is a lattice in cell *σ*, *a*_*σ*_ is the area for the cell *σ* and 𝜁 is a coefficient that constrains single cell *s*(*σ*, *t*).

### Secretome induces protrusion force regulator R

Protrusion force is the driving force for tumor migration (41–43). In many studies, researchers define migration driving force as retraction and protrusion force separately. In our model, we did not explicitly distinguish them. There are no limitations to the polarity of the protrusion force, as we did not differentiate between the front and rear of the cell at the single-cell scale. The direction of the protrusion force can either cause retraction when pointing inwards to the tumor center or promote protrusion when pointing outwards. We have shown that the conditioned media (CM) from hypoxic microtumors induced migratory phenotype in the non-migratory 150µm microtumors (21). To mathematically model this behavior, we hypothesize that conditioned media from hypoxic tumors induce a to-be-identified protrusion force regulator, R, capable of mediating cell protrusion. In our multi-scale microtumor model, the production rate of the protrusion force regulator R is composed of a ubiquitous basal rate *b* and a secretome signal-induced activation rate *s*, where *s* is a fraction of uptake of secretome signal at single cell level, calculated from Eq. 8. In this way, the tumor cells at the boundary of the tumor in contact with the media receive more secretome signals than the cancer cells in the center of the tumor, and thus, will have higher R at the boundary. The spatial distribution of R is represented by the secretome-induced activation rate *s*. The R dynamics can be described as follows:

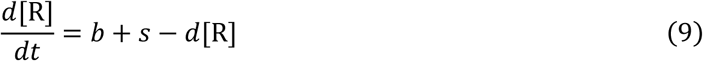

Where *b* is a basal production rate of R, independent of secretome signals in the microenvironment, with the unit of *μM*/*s*. The term *s* is the secretome signal-dependent [*R*] production rate, calculated as *s*(*σ*, *t*_*n*_) at single cell level in Eq. 8. The *d* is the degradation constant for [*R*] with the unit *s*^−1^. Solving for mean field steady state, we have 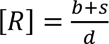. The model parameters can be found in **Supplementary Table S2**. **Protrusion force induced by the regulator R**

Consider all cells can form protrusions with different protrusion forces. Our tumor migration model involves two main factors contributing to the migration modes: the protrusion force and cell-cell adhesion (4,44). Here, regulator R is capable of generating or regulating protrusion force. Possible force-generating components may include cell cytoskeleton, such as F-actin and myosin (45), and Rho signaling family proteins, such as Rac1 (28). Thus, protrusion force F⃑ _*σ*_ for cell *σ* is described as follows, where *f* is the maximum protrusion force.

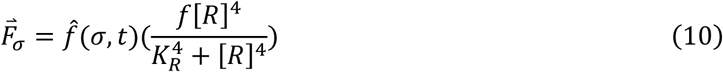

The impact of [*R*] on cellular protrusion force is described as a Hill function, with half saturation concentration *K*_*R*_. Higher [*R*] can promote stronger protrusion for single cells. We denoted the direction of protrusion force F⃑ _*σ*_ acting on cell *σ* as the force polarity vector *p̂*_*m*_(*σ*, t), a unit vector.

In our model, the cell polarity is represented as the direction of cell movement. The direction of protrusion force *f̂*(*σ*, t) for cell *σ*, is determined by the cell’s own motion direction *p̂*_*m*_(*σ*, t) and neighboring cells’ motion direction. The cellular motion direction, i.e., cell polarity vector, *p̂*_*m*_(*σ*, t) is defined as a normalized positional displacement vector calculated from the past 5 MCS (Monte Carlo Steps), representing the cell’s direction of motion, as shown below.

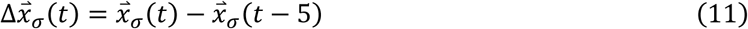

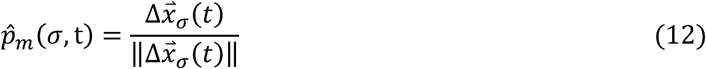

Where *x⃑*_*σ*_(*t*) is the cell *σ*’s position vector at time *t*. Δ*x⃑*_*σ*_(*t*) represents the cell displacement vector in the past 5 MCS. While cellular local alignment is linked to cell-cell adhesion, single cell polarity *p̂*_*m*_(*σ*, t) is independent of local alignment. Local polarity dissipation happens as friction exists between neighboring cells, where the direction of neighboring cell motion impacts the direction of motion of the cell *σ* (46,47). The force direction *p⃑*_*f*_(*σ*, t) is a unit vector defined by the following expression:

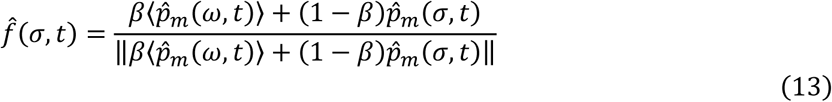

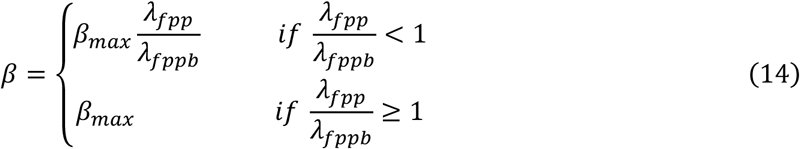

In Eq. 13, *ω* represents cell identifiers that are neighbors to cell *σ*. The force direction *f̂*(*σ*, t) is defined as a linear combination of averaged neighboring cell polarity and the cell’s own polarity. *β* represents the coupling strength of local cellular polarity alignment, which is the fraction of cellular polarity imposed by neighboring cells due to cell-cell interaction, with *β* ∈ [0, *β*_*max*_]. *β* is coupled with adhesion constraint *λ*_*fpp*_ in the model, as shown in Eq. 13. The second term (1 − *β*) in Eq. 13 is the fraction of cell *σ*’s motion polarity *p̂*_*m*_(*σ*, t) contributing to the direction of protrusion force. This term indicates that the cell’s polarity persisted for a certain period.

For Eq. 14, *β*_*max*_ is the maximum possible fraction of polarity that neighbors can impact a cell. *λ*_*fppb*_ is a constant for basal adhesion strength, and *λ*_*fpp*_ is the adhesion strength defined in Eq. 4. Eq. 14 describes the coupling effect that stronger cell-cell adhesion allows neighboring cells to impose a more substantial polarity alignment on cell *σ*.

We also examined cases where Eq. 14 constraints were removed using *β* as a free parameter (Supplemental Materials and Figure S4), recognizing local alignment of cellular polarity as crucial for the emergence of directional migration mode.

### Parameter scan for max protrusion force and adhesion strength

Two key parameters that determine tumor migration mode were scanned, including the maximum protrusion force (*f*) generated by a cell and the strength of adhesion interactions between neighboring cells (*λ*_*fpp*_). For each set of parameters, the simulation was run for 390 Monte Carlo steps (MCS) and repeated 50 times. Each MCS represents 24 minutes in real-time. This was determined by running the simulation until a microtumor morphology comparable to the 150CM tumors in culture for six days was achieved (note: the secretome was added on day 3). We found that a 180 MCS simulation corresponds to 3 days of microtumor development well, with each MCS representing 24 minutes in real-time.

The simulation included three steps. The first step was to relax the model to remove the dependence on the initial state set for the simulation. At the beginning of the simulation, the model was relaxed for 120 MCS. We simulated the initial non-migratory tumor phenotype in the second step without any secretome signal. This step was run for 90 MCS with basal [R] production rate *b* = 1 *μM*/*s* and secretome signal-dependent [*R*] production rate, *s* = 0 *μM*/*s* (no secretome, initially), mimicking the non-migratory 150 μm microtumor condition. In the third step, we simulated a 150μm microtumor treated with the hypoxic conditional media (secretome), setting the maximum secretome signal at the tumor boundary to *s* = 4 *μM*/*s*, with *b* = 1 *μM*/*s*. After adding secretome, the simulation was run for 180 MCS, corresponding to 3 days of the experiment. For each simulation, we collected time-lapse information, such as single cell positions, [*R*] level, force F⃑ _*σ*_, locally transmitted polarity fraction *β*, cell center of mass position. With the collected data, we obtained different migration modes. A phase graph can be found in **Figure S1**.

Total simulation time is a factor that may cause the migration landscape to change. For example, with a longer simulation time, the fraction of directional migration is expected to increase, as observed experimentally.

### Identification of tumor migration modes

The migration modes of simulation results were determined with a two-step unsupervised clustering procedure using a set of nine geometric features of the microtumors. **Figure S2** shows the distribution for all 19 features. First, the migratory and non-migratory modes were identified with six distance-related measures. Then, the directionally and radially migratory tumors were classified by distance and geometric shape features.

In step 1, to identify the migratory modes and non-migratory modes, we performed k-means clustering on the six distance-related features, including the displacement of the center of the tumor mass *D*_*tumor*_, average cell displacement from initial tumor center *d̅*, the standard deviation for cell displacement *D*_*σstd*_, the average of radial distance measure to initial tumor center *D*_*radialavg*_, the standard deviation of radial distance measure to initial tumor center *D*_*radialstd*_, and the standard deviation of cell migration distance comparing to the initial condition *D*_*migrationstd*_. Definitions of the distance measures are given in **Supplementary Table S3**. Features were preprocessed before clustering to avoid bias and optimize clustering results. First, the distance features of the final time frame (270 MCS) were divided by initial condition values (except for the *D*_*migrationstd*_), then normalized to a standard distribution with mean zero and standard deviation 1. Then, the normalized features were clustered with k-means and clustered into two groups corresponding to migratory and non-migratory tumors. **Figure S2c** gives the clustering result in the uniform manifold approximation (UMAP) space.

In step 2, to identify the radial and directional migratory tumors, we used a total of 19 geometric features, including eight distance features and 11 shape features (48). **Figure S2a** shows the distribution of the 19 geometric features of the simulated tumors. Again, we preprocessed the features before clustering. The features were first divided by their initial step values and then normalized to a standard distribution with a mean of 0 and a standard deviation of 1. Next, we performed feature dimensional reduction with UMAP, followed by k-mean clustering on the reduced features. We used a fixed random seed for both UMAP and k-means for reproducibility. **Figure S2d** gives the clustering results in the UMAP space.

## Results

### 1. Tumor-intrinsic hypoxia induces directional migration in large microtumors, while hypoxic secretome induces radial migration in small microtumors

We have previously fabricated hydrogel microwell arrays (**Figure 1a**) for studying microtumor migration under controlled microenvironments (18,23). The uniform-sized microtumors are generated from the same non-invasive parental T47D breast cancer cells using microwells either with a diameter of 600 µm (denoted as large 600 microtumors) or with a diameter of 150 µm (denoted as 150 microtumors). Our prior studies indicated that the large 600D6 microtumors displayed migratory behavior, whereas the small 150D6 microtumors remained non-migratory for the entire culture time of six days (where D6 indicates six days of culture time) (19–21,23). Large 600 microtumors spontaneously developed tumor-intrinsic hypoxia as early as day 1, started directional collective movement of the whole tumor cluster from day 3, and upregulated expression of mesenchymal markers without loss of E-cadherin (19,21,23). This directional collective migration is characterized by an initial movement of the tumor towards one side of the microwell and eventually “migrating out” of the microwell (**Figure 1b**) (19,20,23). We further treated the non-migratory, non-hypoxic small 150 microtumors with conditioned medium (CM, hypoxic secretome) of the migratory large microtumors starting from day 3 (denoted as 150CM microtumors). The secretome treatment induced a distinct migratory phenotype in 150CM microtumors, where the cells migrated outward radially from the center of the microtumor (**Figure 1c**). The secretome-induced radial migration is characterized by the appearance of multiple migratory fronts in random directions.

To understand the biophysical mechanisms for the emergence of such distinct tumor migration modes, we performed RNA microarray analysis for directionally migrating 600D6 tumors and radially migrating 150CM tumors, using non-migratory 150D6 as a reference (23). We focused on the differentially expressed genes, which were further enriched for Gene Ontology (GO) terms for biological processes (BP). GO terms describing biophysical processes during tumor migration include “Actin filament organization”, “Biological adhesion”, “Regulation of cytoskeleton organization”, and so on (**Figure 1d**). Microarray analysis of hypoxia-induced directional migration (600D6) revealed that cell adhesion-related terms such as “Cell-cell adhesion”, “Biological adhesion”, and “Positive regulation of locomotion” were the top positively enriched BP with the highest normalized enrichment scores (NES). The GO term “Biological adhesion” included 131 genes differentially expressed in directionally migrating 600D6 compared to non-migratory 150D6. In the analysis of secretome-induced radial migration, *i.e.,* 150CM microtumors, cytoskeletal organization-related terms such as “Positive regulation of cytoskeleton organization”, “Actin filament organization” and “Supramolecular fiber organization” were the top positively enriched BP when compared to non-migratory 150D6 microtumors. Adhesion-related processes, such as “Biological adhesion”, “Cell-cell adhesion”, and “Cell junction assembly” were downregulated in 150CM microtumors in contrast to 600D6. On the other hand, protrusion force-related processes, such as “Actin filament-based process”, “Microtubule polymerization or depolymerization”, and “Regulation of cytoskeleton organization” were downregulated in directional migration (600D6) and upregulated in radial migration (150CM). Together, GO term enrichment analyses suggest a role of cell adhesion in directional migration phenotype, while actin-microtubule cytoskeleton reorganization may be necessary for secretome-induced radial migration phenotype.

### 2. A multi-scale microtumor model (MSMM) incorporates intercellular dynamics and intracellular polarity alignment

#### 2.1 An overview of the MSMM model

Based on the RNA microarray analysis results, we deduced that mechanistically differential regulation of the protrusion force and adhesion plays a crucial role in the emergence of different tumor migration modes. To understand the emergence of diverse collective migration modes, we hypothesize that strong protrusion force in single cells is critical for microtumor migration, while strong cell-cell adhesion aids directional collective migration. To test our hypothesis, we built a tumor migration model based on the **m**ulti-**s**cale **m**icrotumor **m**odel (MSMM), which is based on the cellular Potts model (CPM) on a 2D plane. CPM is a type of agent-based modeling. It represents cells on a lattice, where each cell occupies a set of lattice sites. The model simulates cellular behaviors by minimizing an energy function, which allows straightforward implementations of various cellular activities, such as motility, adhesion, proliferation, and apoptosis, while keeping a realistic cellular shape (45–49). We modeled small 150CM microtumors where the tumor-intrinsic hypoxia stress is not severe (21,23). An illustration of the model regime is shown in **Figure 2**.

**Figure 2.**
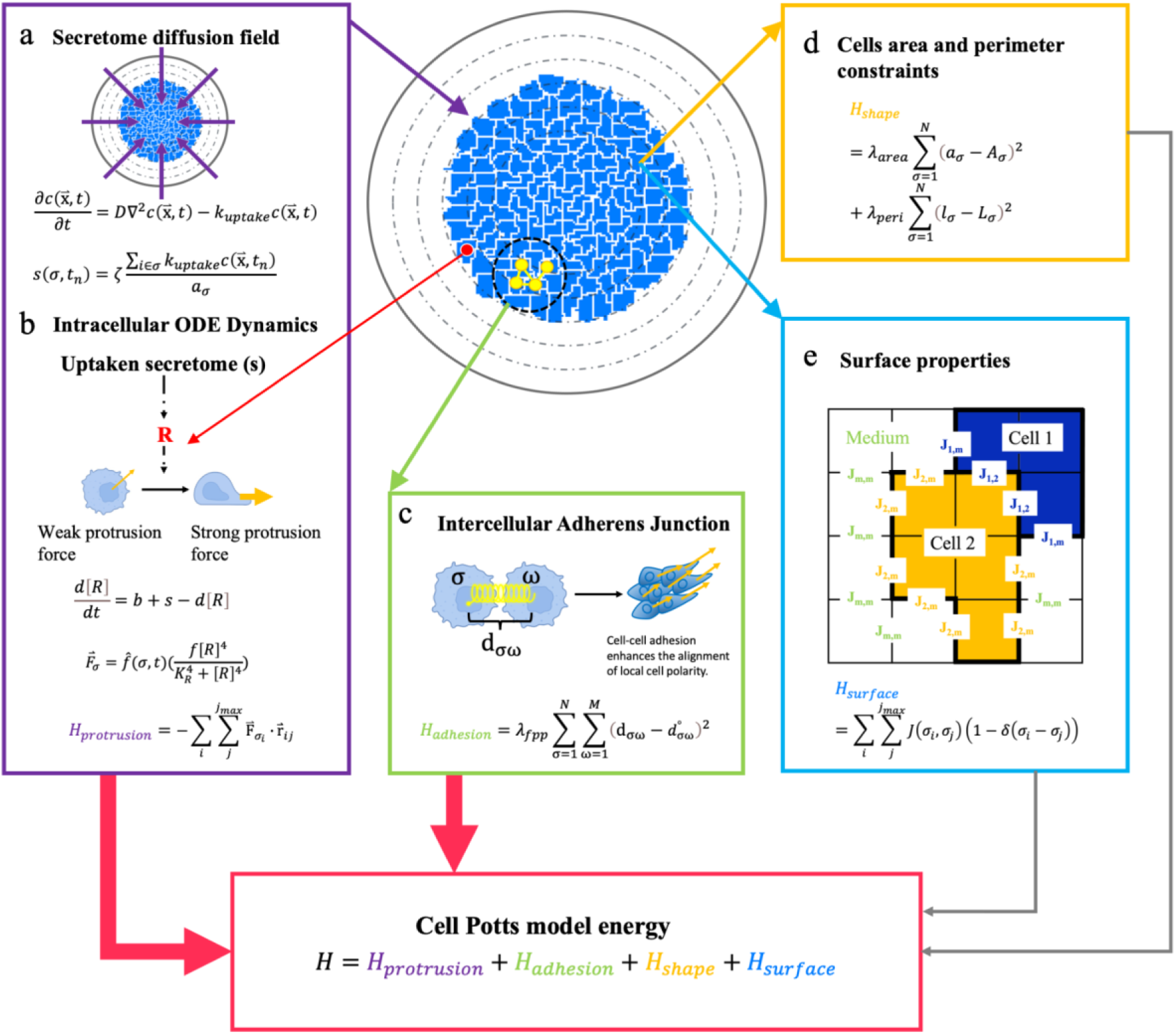
A multi-scale microtumor model (MSMM) scheme consists of four energy terms: energy from the protrusion force, adhesion energy, shape constraint energy, and surface energy. Tumor cells are colored in blue on the lattice representation, while the secretome signal diffuses into the tumor and is represented as concentric circles. The stochastic multi-scale microtumor model simulation was performed by accepting an energy-favorable random move for lattice copy attempts (Eq. 6). **a.** At lattice position x⃑ with a local secretome signal concentration *c*(x⃑, *t*) at time t, a cell (*σ*) can uptake the secretome signal with an uptake rate constant *k*_*uptake*_. Equation details can be found in Eq. 7-8. **b.** The averaged secretome signal taken up by each cell (*s*) induces the production of [R] and subsequently generates protrusion force. Details can be found in Eq. 5, 9 and 10. **c.** Cell-cell adhesion is modeled as elastic strings linking the centers of mass for pairs of cells within a defined distance, with a corresponding energy term as a quadratic term for the distance between cells (d_σω_) with respect to a target distance (*d*^°^) (Eq. 4). In the model, we coupled cell-cell adhesion and local alignment of cellular polarity, as described in Eq. 13-14. **d.** For each cell, the cell shape energy *H*_*s*ℎ*ape*_ is constrained with quadratic energy terms for cell area (*a*_*σ*_) and perimeter (*l*_*σ*_), where *A*_*σ*_ and *L*_*σ*_ are the target area and perimeter corresponding to *a*_*σ*_ and *l*_*σ*_ (Eq. 2). **e.** Lattice representation for surface energy *H*_*surface*_ between different interfaces, such as cell-cell (*J*_1,2_), cell-medium (*J*_1,*m*_ and *J*_2,*m*_) and medium-medium (*J*_*m*,*m*_) (Eq. 3), are shown.

For a single cell in a microtumor, protrusion force can be generated by localized actin polymerization at the plasma membrane, and cell-cell adhesion is regulated by adherens junction (44). For 150 microtumors, the secretome plays key roles in inducing tumor migration. In the MSMM, cells take up the secretome factors from conditioned media (CM) at a rate depending on their position in the microtumor. For example, cells at the periphery have a higher uptake of the secretome than those in the core of the microtumor because less secretome is inside the tumor than at the boundary of the tumor. We simplify the secretome factors as a continuous field that follows reaction-diffusion dynamics (**Figure 2a**). We assume that the secretome factors up-regulate intracellular protrusion force regulator R, a protein that can promote lamellipodia formation and mediate protrusion force, leading to increased protrusion force (**Figure 2b**). Potential candidates for the protrusion force regulator may include Rac1, one of the significant signaling proteins that regulate cytoskeletal rearrangements (49) and myosin II, a motor protein that controls the force exerted by lamellipodia and filopodia (50). In MSMM, a cell can protrude in random directions, *i.e.,* protrusion attempts toward the tumor periphery or center.

Cell-cell adhesion preserves tissue integrity by passively resisting forces that can act on tissues and cells (51). Different cell junctions, such as adherens junctions, tight junctions, and desmosomes, provide tissue adhesion and mechanical support and facilitate signal transduction between cells (52). Some cell-cell adhesion molecules have been implicated in the regulation of cell polarity because cell-cell adhesion molecules directly interact with or regulate the polarity molecules and pathways (53). Neighboring cells, as a key extrinsic factor, are observed to regulate cell polarization. For example, E-cadherin is a major cell-cell adhesion molecule. E-cadherin-mediated cell-cell adhesion can activate Cdc42, an essential protein regulating cell polarity (54). Inhibiting E-cadherin junctions blocks proper polarization and produces blastocysts (55,56). *CRB3* encodes for Crumbs cell polarity complex component 3 and is crucial for cell polarity in cancer development (57,58). E-cadherin expression level is also linearly correlated with *CRB3* expression level in cancers, with a Pearson correlation of 0.81, as shown in **Figure S3** (59). Moreover, neighboring cells mechanically induce traction forces and shear stress on a cell, affecting its polarity (60,61). In our model, we describe an “adhesion energy” term, which incorporates intercellular, membrane, and cytoskeletal forces acting on the cell-cell adhesion interface (51) (**Figure 2c**). The cell-cell adhesion strength is then coupled with cellular polarity local alignment. Detailed model descriptions on cell-cell adhesion coupling with local alignment of cellular polarity can be found in **Materials and Methods** and **Eq. 11-14**. Further discussion on how independent local alignment variables impact model dynamics can be found in Supplemental Materials and **Figures S3-S4**.

Each cell can occupy multiple lattice sites, and cell area and perimeter are constrained by energy terms (**Figure 2d-e, Eq. 1-3**). Together, the model incorporates intercellular dynamics and intracellular polarity transmission through tension. We implement our model with the open-source software CompuCell3D (CC3D) (33), the details of which are described in Methods. The model and analysis code can be found at https://doi.org/10.5281/zenodo.14928214

#### 2.2 The multiscale microtumor model recapitulates distinct migratory modes

To recapitulate the distinct microtumor migration modes induced by tumor-intrinsic hypoxia or secretome (21), we used the multiscale microtumor model to simulate the induction of protrusion force regulator R and cell-cell adhesion in response to the secretome signal (**Figure 3**). Each simulation corresponds to roughly six days of cell culture time to mimic our experimental setup of 3D microtumor migration. Details of the mathematical modeling are in the **supplementary** section, and the parameters used in this model are described in **Tables S1 and S2**.

**Figure 3.**
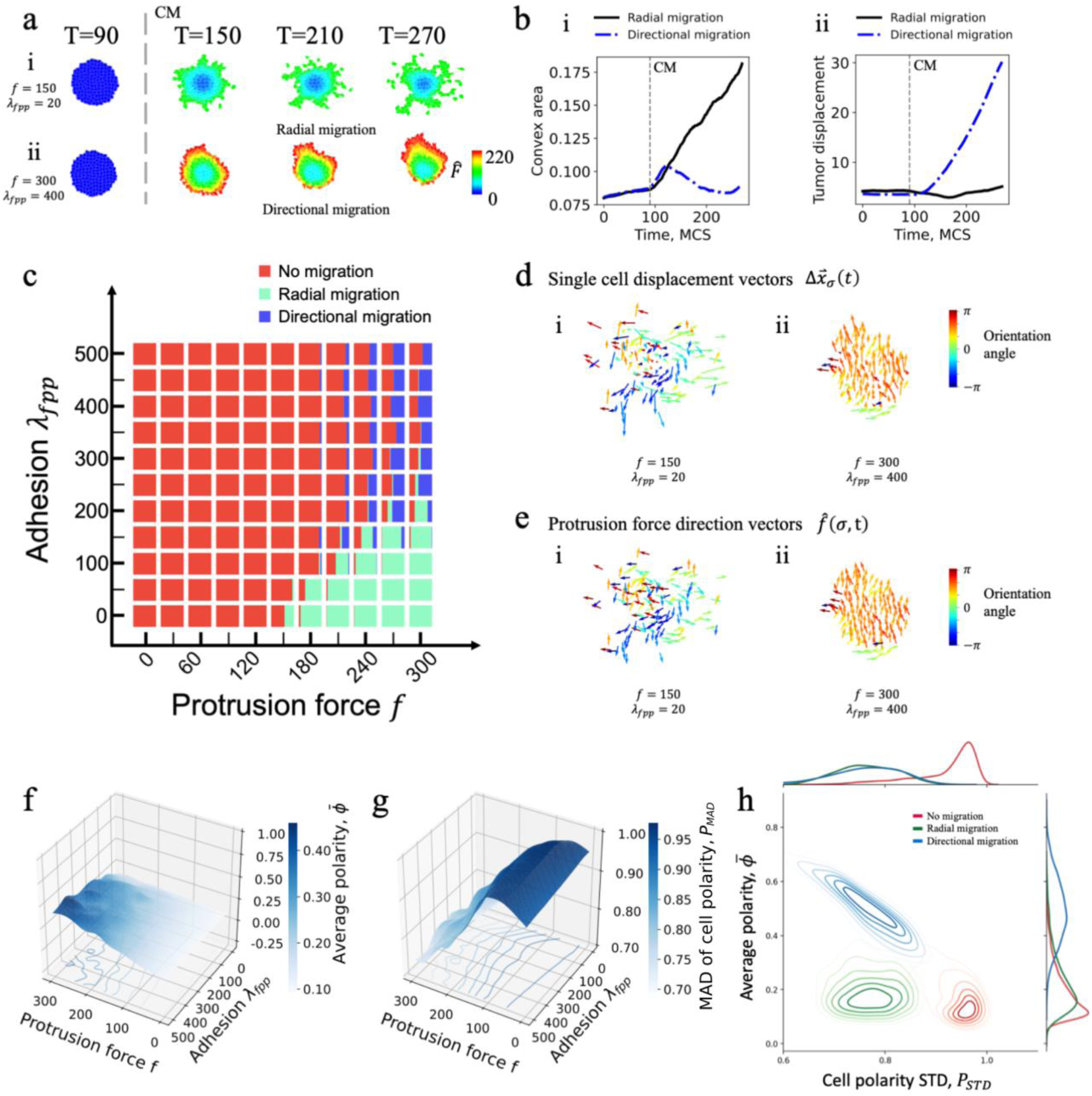
The multiscale microtumor model recapitulates distinct migration modes. **a.** Representative snapshots showing (i) radial migration with *f* = 150 and *λ*_*fpp*_ = 20 and (ii) directional migration with *f* = 300 and *λ*_*fpp*_ = 400. **b.** Temporal trajectories for (i) convex hull area and (ii) tumor displacement corresponding to those in panel a. The x-axis is time in Monte Carlo steps (MCS). **c.** Fractions of the tumors showing different migration modes in the parameter space of protrusion force *f* and adhesion strength *λ*_*fpp*_. **d.** Representative snapshots of the cell displacement vector Δ*x⃑*_*σ*_(*t*) for the past 5 MCS for (i) radial migration and (ii) directional migration for two simulations in panel a. e. Representative protrusion force direction *f̂*(*σ*, t) snapshots for (i) radial migration and (ii) directional migration. **f.** Average tumor polarity *ϕ̅* shows high average polarity for high protrusion force *f* and high adhesion *λ*_*fpp*_. **g.** The mean absolute temporal deviation of cell polarity *P*_*MAD*_ decreases as protrusion force *f* increases, while adhesion strength *λ*_*fpp*_ has less impact on *P*_*MAD*_ compared to the protrusion force *f*. **h.** Gaussian kernel density estimation (KDE) plot for distribution of tumor migration modes in the parameter space of cell polarity variance *P*_*MAD*_ and tumor average polarity *ϕ̅*. The levels for kernel density are [0.2, 0.4, 0.6, 0.8].

To better illustrate the interaction between protrusion force and adhesion, we defined an effective force *F̂* to quantify the real-time protrusion force distribution across cells within a tumor:

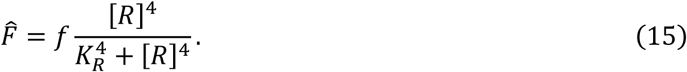

The impact of [*R*] on cellular protrusion force is described as a Hill function, with half saturation concentration *K*_*R*_. Higher [*R*] can promote stronger protrusion for single cells. Our simulation results reveal distinct microtumor migration behaviors depending on the model parameters. Specifically, two key parameters that determine tumor migration mode are the maximum protrusion force (*f*) generated by a cell and the strength of adhesion interactions between neighboring cells (*λ*_*fpp*_), for example, with *f* = 150 and *λ*_*fpp*_ = 20 (both in arbitrary units), a tumor migrates radially with cells migrating in different directions (**Figure 3a**). With *f* = 300 and *λ*_*fpp*_ = 400, the tumor migrates directionally, where cells collectively migrate as a cluster in one direction. When individual cells migrate radially from the microtumor, the overall convex area of the migrating microtumor increases while the whole tumor displacement remains low. In contrast, the directionally migrating microtumor displays a low convex area and an increase in the whole tumor displacement over time (**Figure 3b**).

To systematically investigate the parameter dependence, we performed a parameter scan over the maximum protrusion force within a range of [0,300] and adhesion strength within a range of [0,500]. We characterized the simulated tumors using 19 geometric features (**Supplementary Table S3** and **Supplementary methods** for details), which cluster the simulated tumors into non-migrating, radially migrating, and directionally migrating tumors (**Figure S3b-c**).

Next, we investigated how the distribution of migration mode changes with values of the maximum protrusion force *f* and adhesion *λ*_*fpp*_ (**Figure 3c**). The radial migration mode is dominant in a region with high protrusion force and low adhesion, while the directional migration mode is dominant in a region with high protrusion force and high adhesion. No migration was observed when the maximum protrusion force *f* is low (*f* < 150). These results suggest that a certain threshold of protrusion force is necessary for inducing migration. Above this protrusion force threshold, the cell-cell adhesion strength dictates whether tumors assume directional or radial migration mode.

In radially migratory tumors, directions of cell migration are random, while in directional migration, directions of cell migration are more aligned, as shown in *Figure 3d-e*. Cell polarity (*p̂*_*m*_(*σ*, t)) is represented as the direction of cell movement (Eq. 12). In our model, cell-cell adhesion strength is coupled with local cellular polarity alignment (Eq. 13-14). To systematically quantify the correlations between tumor cell polarities during the migration, we defined a normalized average cell polarity in a microtumor as an order parameter (62),

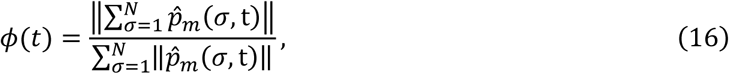

Where polarity *ϕ*(*t*) describes the alignment of migration directions of various cells within the tumor at time *t*, and it varies between 0 and 1. When *ϕ*(*t*) = 0, the tumor cells move in random directions. When *ϕ*(*t*) = 1, the cell migration directions are completely aligned.

The secretome (conditioned medium, CM) activates peripheral cells to generate protrusion forces that drive the tumor into migratory states. To facilitate the characterization of the dependence of tumor polarity on cell adhesion strength, we defined a temporally averaged tumor polarity *ϕ̅* as

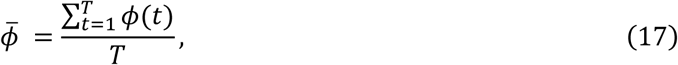

which describes the alignment of migration directions of all cells within the tumor during the simulation time interval *T*. It is an order parameter ranging between 0 to 1. When *ϕ̅* = 0, the directions of cell migration within a tumor are random, and the tumor likely migrates radially. When *ϕ̅* = 1, the migration directions of all cells within a tumor are aligned, where the tumor likely migrates directionally. Furthermore, we defined the mean absolute temporal deviation of cell polarity as an order parameter *P*_*MAD*_,

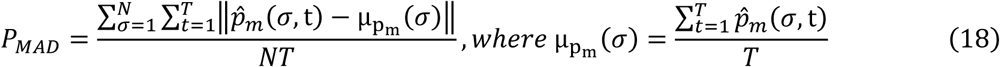

The *p̂*_*m*_(*σ*, t) is the polarity vector for cell *σ* at time *t*, and μ_pm_(*σ*) represents the temporal average cellular polarity for cell *σ* over the simulation time T. The order parameter *P*_*MAD*_ describes the randomness of the cellular migration directions over the simulation period in a tumor. Our results suggest that different migration modes show distinct polarity properties (**Figure 3f-h**). In non-migratory tumors, the average polarity *ϕ̅* is low, while the cell migration directions are random, with higher *P*_*MAD*_ compared to radial and directional migration. Radially migrating tumors are less polarized (low *ϕ̅*), with cellular polarity over time (*P*_*MAD*_) being less random compared to non-migratory tumors (**Figure 3h**). This leads to multiple migratory fronts in radial migration, where different groups of cells choose to migrate in different directions. Compared to radial migration, directionally migrating tumors have higher average polarity (high *ϕ̅*) but similar cell polarity noise over time (low *P*_*MAD*_), thus the cells are better aligned in one direction.

In summary, parameter scan simulations reveal that cellular protrusion force and cell-cell adhesion promoted local cell polarity alignment are crucial determinants of distinct migration patterns. First, cells need to exert enough protrusive force to initiate tumor migration. Radial migration manifests when migrating cells align locally and move collectively in diverse directions. Directional migration, on the other hand, results from symmetry breaking. With increased cell-cell adhesion strength, more substantial polarity alignment corresponds to a strong alignment of protrusion force in one direction that overcomes protrusion forces in other directions, resulting in the entire tumor cluster moving in one direction. Using the present model, we show that the spatial correlation length increases with *λ*_*fpp*_ (**Figure S5**).

Taken together, a sufficient protrusion force is necessary for any kind of migration, while strong cell-cell adhesion is necessary for directional migration.

### 3. Reducing protrusion force attenuates radial tumor migration

We next sought to understand the impact of protrusion force on tumor migration. In MSMM, force *f* exerted in single cells is regulated by a general force regulator [*R*], under the influence of the secretome. We examined the distribution of tumor migration modes by varying maximum protrusion force *f*, at constant cell-cell adhesion strength at *λ*_*fpp*_ = 50 **(Figure 4a)**. With this relatively low cell-cell adhesion strength, tumors either remain non-migratory or exhibit radial migration, recapitulating our experimental observation of radially migrating 150CM microtumors after CM treatment (21,23). With an increasing protrusion force, the dominant migration mode changed from non-migratory to radial migration (**Figure 4a**), where the percentage of radially migrating tumors increased with increasing protrusion force. For example, with a stronger protrusion force *f* = 200, the tumor has a strong radial migration phenotype, showing strands extending in all random directions (**Figure 4b**). The convex area of microtumors, *A_tumor_* (normalized by the total simulation area, **Supplementary Table S3**), and cell displacement distance, *d̅* (measured from the initial tumor center, **Supplementary Table S3**) both increased with stronger protrusion force (**Figure 4c and 4d**).

**Figure 4.**
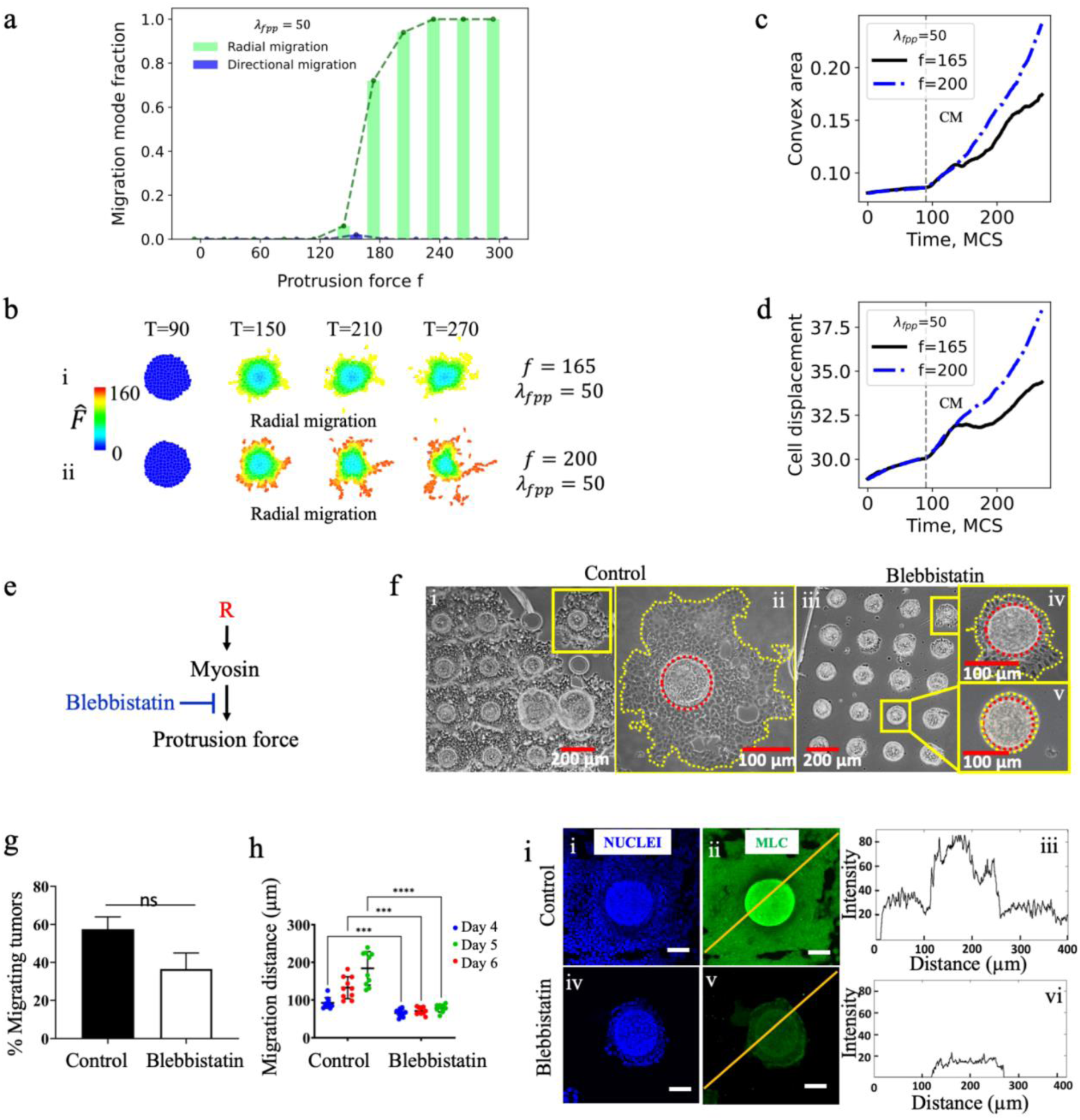
Reducing protrusion force attenuates radial tumor migration. **a.** Distribution of fraction of tumor migration modes as a function of varying protrusion force in the parameter scan with constant adhesion strength *λ*_*fpp*_ = 50. **b.** Typical time trajectories for simulation with (i) weak (*f* = 165) and (ii) strong (*f* = 200) protrusion force at a constant adhesion strength of *λ*_*fpp*_ = 50. The single cell protrusion force *F̂* is color-coded. The time trajectories for **c.** convex area and **d.** average cell displacement from the initial center of the tumor with weak (black line) and strong (blue) protrusion forces are shown. The x-axis is time in Monte Carlo steps (MCS). **e.** Inhibition of myosin II by blebbistatin impairs the generation of protrusion force. Panels **f-h** show the effect of myosin inhibition on 150CM tumors using blebbistatin treatment. f. Brightfield images showing significantly reduced migration distance in (iii-v) blebbistatin-treated 150CM tumors on day 6 compared to (i-ii) untreated control. **g.** Percentages of migrating tumors did not decrease significantly on day 6 in the blebbistatin-treated group compared to the control (*p-value* = 0.0755; paired t-test; *n* = 2 devices, each containing 484 microtumors from two independent biological experiments). **h.** The extent of migration was significantly reduced in 150CM microtumors in blebbistatin-treated microtumors compared to the untreated control. (*p-value* <0.0005, two-way ANOVA, *n* = 10 representative microtumors, N = 2 devices). **i.** Confocal microscopy images of control and blebbistatin-treated microtumors after immunostaining with Hoechst (nuclei; blue) and myosin light chain (MLC; green) antibody. The myosin light chain (MLC) expression is reduced in the blebbistatin-treated microtumor (iv-v) compared to the untreated control (i-ii). The respective intensity plot profiles indicate the decreased intensity of MLC and reduced extent of migration in the blebbistatin-treated microtumors compared to the control.

Many factors can affect the protrusion force. Both pushing (protrusive) and pulling (contractile) forces are critical for cell migration (45,63). A pushing force generated by coordinated polymerization of actin filaments can be regulated by Rac1, amongst many other factors. On the other hand, a pulling force can be generated by sliding actin filaments along myosin II filaments (45). Myosin II, a key effector for Rho kinase signaling (64), can promote contractility and generate protrusion force by forming actomyosin filaments (65–67). Abhinava *et al.* reported that myosin II regulates protrusion dynamics and can convert border cells into migratory cells in collective migration (43). The functionality of myosin II has been extensively studied *in vitro* and *in vivo*. Therefore, we used blebbistatin to inhibit myosin II (**Figure 4e**) experimentally.

When treated with blebbistatin, 150CM microtumors showed a significant reduction in the extent of migration (**Figure 4f-h**). Since we counted all the tumors that migrated out of the microwells as ‘radially migrating’ tumors irrespective of their extent of migration, we did not observe a significant reduction in the percentage of migrating tumors in the blebbistatin-treated group (36.5± 14.6%) compared to the untreated 150CM microtumors (57.5 ± 11.2%) (p value=0.0755; paired t-test; n = 4 devices each containing approximately 480 microtumors from two independent biological experiments) (**Figure 4g**). However, distance of tumor migration showed a significant decrease in blebbistatin-treated tumors, while untreated 150CM tumors continued to migrate out of the microwell from day 4 to day 6. Indeed, the distance of migration reduced from 184 ± 44 µm for untreated 150CM microtumors to 78 ± 10 µm for blebbistatin-treated 150CM microtumors on day 6 (**Figure 4h**), suggesting inhibition of migration due to myosin II inhibition. This observation was further confirmed by reduced myosin light chain (MLC) expression in blebbistatin-treated microtumors compared to untreated control (**Figure 4i**). Our experimental results confirmed our model prediction that reduced protrusion force reduces radial migration.

### 4. Increased cell-cell adhesion decreases radial migration while promoting directional migration

Directional migration was first observed in large microtumors under tumor-intrinsic hypoxia conditions. Our RNA microarray analysis revealed key directional migration gene signatures, including PIK3R3 (23), a regulatory component for PI3K. Interestingly, pathway enrichment analysis revealed that the PI3K-AKT pathway is differentially regulated in directional and radial migration modes. AKT, a serine/threonine protein kinase, plays a crucial role in the PI3K/AKT pathway, influencing cell metabolism, growth, proliferation, and survival (68). PI3K activates AKT [3], which comprises three isoforms: AKT1, AKT2, and AKT3 (69). Several studies have established that the isoforms AKT1 and AKT2, play distinct and opposing roles in cell signaling and tumor growth and progression (70–72). A study by Polytarchou *et al.* (73) further elucidated the association between AKT and hypoxia and reported that only AKT2 is induced under hypoxic conditions, not AKT1 or AKT3. Also, AKT2-expressing cells were more resistant to hypoxia-mediated cell death and hypoxia-induced cell cycle G1/S arrest. Furthermore, under oxygen deprivation, AKT2, via PTEN inhibition, acts as a master regulator of all three isoforms of AKT, ensuring higher cell survival under hypoxia (73). Among AKT isoforms, our microarray analysis revealed that AKT2 is differentially regulated in hypoxia-induced directional migration and secretome-induced radial migration (23). Since the role of AKT2 in 3D collective migration remains understudied, we further studied the effect of AKT2 inhibition on 150CM microtumor migration.

Upon treatment with CCT128930, a potent AKT2 inhibitor, the percentage of migratory microtumors (16.4 ± 3.9) reduced significantly (*p* value= 0.01) compared to untreated 150CM group (40.1 ± 12.6) (**Figure 5a-b**). Note that we counted all the tumors that migrated out of the microwells as ‘migrating’ tumors irrespective of their extent of migration. For the migratory tumor population in the CCT128930-treated group, we did not observe significant changes in the extent of migration except on day 6 compared to the untreated control (**Figure 5c**). Within the CCT128930-treated group, we observed three different phenotypes: a) complete inhibition of migration, b) no effect of CCT128930 treatment on inhibition of migration, where 10.9 ± 5.6% of tumors continued to migrate radially, and c) interestingly, the emergence of a directional migration phenotype in 5.5 ± 2.2% microtumors with AKT2 inhibition (**Figure 5a-b**). These data suggest that AKT2 plays a role in determining the collective migration mode. This was further confirmed by the differential expression of pAKT in the radial and directional migration modes, while the expression of total AKT remained unchanged (**Figure 5d**). Our experimental results suggest that AKT2 may play an essential role in the plasticity of collective migration modes of microtumors.

**Figure 5.**
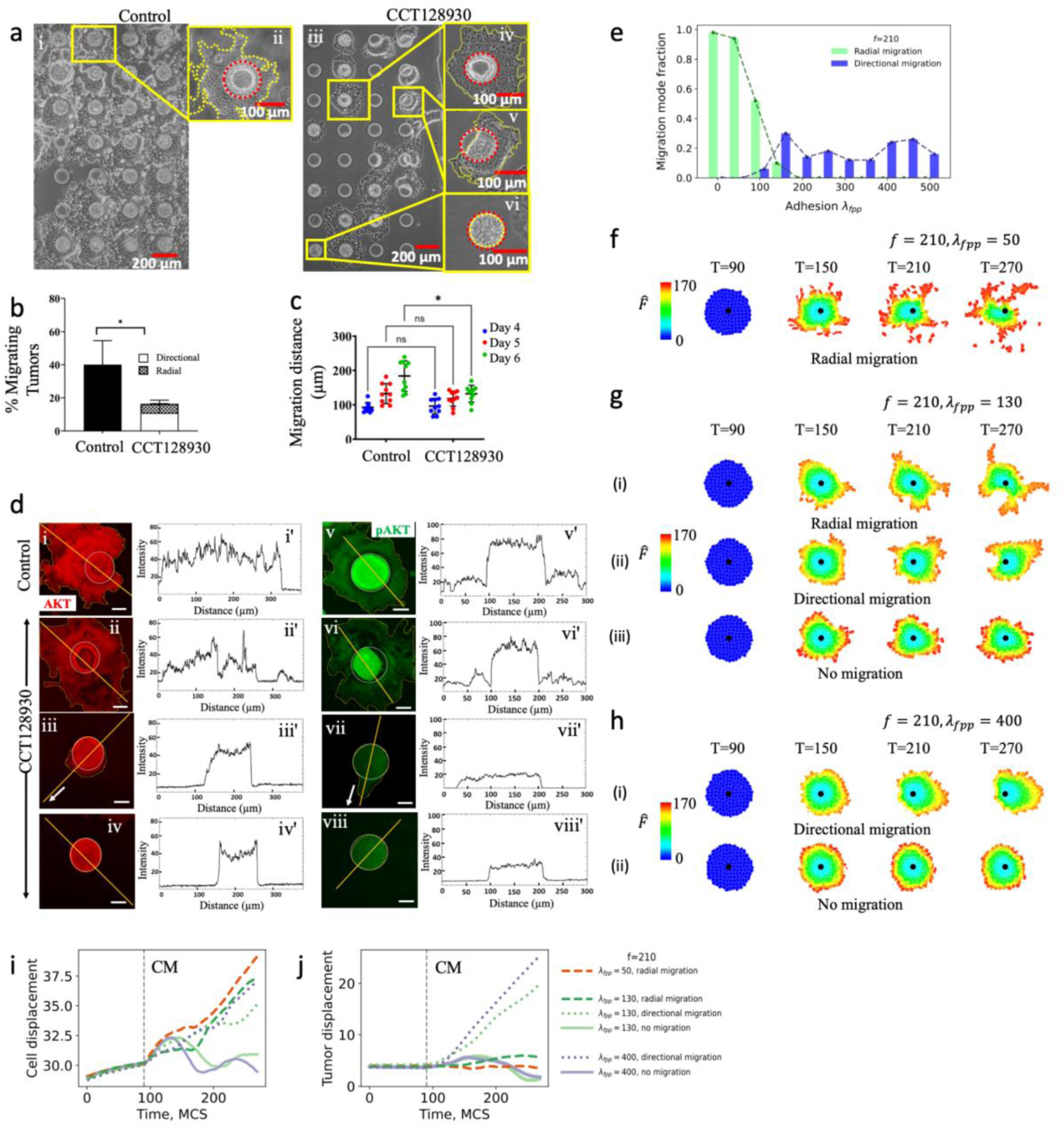
Increased cell-cell adhesion decreases the fraction of radially migrating tumors and promotes directional migration. **a.** Brightfield images showing the dominance of radial migration in untreated controls (i & ii) whereas CCT128930-treated150CM tumors (iii) consist of a mixed population of (iv) radially migrating, (v) directionally migrating, and (vi) non-migratory tumors on day 6. **b.** Percentages of migrating tumors in the CCT128930-treated group decreased significantly on day 6 compared to the untreated control group (‘*’ denotes *p-value* of 0.01; paired t-test, n=2 devices each containing 484 tumors). We observed a mixed population of migrating tumors displaying either radial or directional mode of migration. **c.** Treatment with CCT128930 significantly reduced the extent of migration compared to the untreated control on Day 6 (*p-value* = 0.018, two-way ANOVA, *n =* 10 different microtumors, N = 2 independent experiments). **d.** Confocal microscopy images of control and CCT128930-treated microtumors after immunostaining for total AKT (red) and pAKT (green) (Scale bar = 50µm). Arrows represent the direction of migration. Differential expression of pAKT was observed in CCT128930-treated tumors compared to the control group, while expression levels for total AKT remained unchanged. The respective intensity plot profiles for each image were plotted in ImageJ (i - viii). **e.** At a constant protrusion force *f* = 210, the distribution of tumors in different migration modes (radial versus directional) is dictated by the cell-cell adhesion strength *λ*_*fpp*_. Radial migration mode dominates at low cell-cell adhesion (*λ*_*fpp*_ ≤ 100) values (weak cell-cell adhesion) while strong cell-cell adhesion (*λ*_*fpp*_ > 100) promotes directional migration. The total fraction of radial and directional migratory tumors is also reduced as the cell-cell adhesion strength increases. **f.** Typical simulation time snapshots show radial migration in low adhesion conditions (*λ*_*fpp*_ = 50). Black dots are initial tumor centers. **g.** Under moderate cell-cell adhesion strength *λ*_*fpp*_ = 130, time snapshots for three typical migration modes, (i) radial, (ii) directional, and (iii) non-migratory modes are shown. **h.** Under strong cell-cell adhesion strength *λ*_*fpp*_ = 400, (i) directional migration and (ii) non-migratory modes coexist, and their typical migration time snapshots are shown. **5i** and **5j** show examples of simulated trajectories for cell displacement and tumor displacement under varying cell-cell adhesion strength. The x-axis is time in Monte Carlo steps (MCS).

The cell-cell adhesion plays a central role in cell migration (74). AKT2 may regulate microtumor migration by tuning cell-cell adhesion. For example, cell adhesion pathways are reported to be upregulated in AKT2 knockout cells (69). E-cadherin is a key cell adhesion molecule mediating adherens junction and maintaining the epithelial phenotype of cells (75). Snail1 is a transcriptional factor that is essential for triggering epithelial-to-mesenchymal transition. Snail 1 promote AKT2 kinase binding to *CDH1* promoter, decreasing E-cadherin expression, explaining the pro-mesenchymal role of AKT2 (76,77). The mechanistic role of cell-cell adhesion in the plasticity of tumor migration modes remains elusive. It is also challenging to precisely measure the cell-cell adhesion strength experimentally.

Our MSMM described above shows the critical role of cell-cell adhesion in directional tumor migration. The MSMM model simulations indicate that enhancing cell adhesion strength, along with the coupled local alignment of cellular polarity, results in the emergence of directional migration (**Figure 3c**). Thus, we examined the distribution of tumor migration modes as a function of adhesion strength *λ*_*fpp*_ with maximum protrusion force fixed at *f* = 210 (**Figure 5e**). At a lower adhesion strength *λ*_*fpp*_ ≤ 100, most tumors migrate radially. Increasing adhesion strength *λ*_*fpp*_ leads to a decrease in the fraction of radially migrating tumors. At *λ*_*fpp*_ ≥ 100, directional tumor migration emerges. Thus, the MSMM simulations show that increasing cell-cell adhesion can switch the microtumor migration from radial to directional migration.

**Figure 5f-h** shows example trajectories from simulations with different adhesion strengths. At a low adhesion strength, where *λ*_*fpp*_ = 50, tumors migrate radially. At a moderate adhesion strength, where *λ*_*fpp*_ = 130, two modes of migration, *i.e.*, radial and directional migration modes, co-exist. At a high adhesion strength, where *λ*_*fpp*_ = 400, directional migration mode predominates. Two geometric features, the average cell displacement *d̅* and tumor displacement *D*_*tumor*_ (displacement of tumor centroid, **Supplementary Table S3)** are shown in **Figure 5i-j**. We compared typical time series for cell displacement and tumor displacement under weak (*λ*_*fpp*_ = 50, red line), moderate (*λ*_*fpp*_ = 130, green lines), and strong (*λ*_*fpp*_ = 400, purple lines) adhesion strength conditions, and found that the directional migration has distinct properties compared to the radial migration mode in terms of cell displacement and tumor displacement (**Figure 5i-j**). In **Figure 5i**, after treatment with CM, cell displacement of non-migratory tumors keeps steady at lower values, while the cell displacement keeps increasing in directional and radial migratory tumors. In **Figure 5j**, after treatment with CM, tumor displacement of directionally migrating tumors increases, while the tumor displacement of radial and non-migratory tumors remains steady near zero during the simulation. These observations from the MSMM simulations verified the co-existence of directional and radial migration in moderate adhesion conditions. They also revealed that the directional and radial migration possess distinct properties during the simulation.

To summarize, AKT2 inhibition studies (**Figure 5a-d**) revealed an essential role of AKT2 in regulating the plasticity of microtumor migration modes. At the same time, the MSMM can recapitulate the observed plasticity of collective migration by varying cell adhesion strength. Taken together, we postulate that AKT2 is a critical regulating factor in determining the plasticity of collective migration by regulating adhesion-related biophysical properties.

## Discussion

Collective migration of cells is a hallmark of many physiological and pathological processes, such as embryonic morphogenesis, wound healing, and stage IV cancer progression (66). In cancer metastasis, collective migration confers higher invasive capacity and resistance to therapy compared to single-cell migration (78,79). Clinically, circulating tumor cell (CTC) clusters, prevalent in metastatic patients, exhibit increased resistance to radiotherapy and chemotherapy (78,80–82) and are often associated with worse prognosis (83). Collectively migrating tumor clusters exhibit improved cell survival in various microenvironments, evade the immune system, and colonize secondary sites more efficiently (83). Furthermore, different tumor cells exhibit migratory plasticity, which refers to the ability of tumor cells to reversibly switch between different migration modes, depending on the cellular and microenvironmental context (84–87). Such plasticity adds further complexity to tumor migration mechanisms, necessitating a fundamental perspective model for comprehensive understanding. Here, we employed *in vitro* 3D microtumor models (18–23) and computational modeling to elucidate the mechanisms underlying collective migration in secretome-induced microenvironments. Our multiscale modeling approach integrated single-cell signaling with population-level interactions, focusing on the dynamic interplay between cellular protrusion force and cell-cell adhesion.

Widely studied individual cell migration modes include mesenchymal, which mainly depends on the activity of small GTPase Rac, while amoeboid migration is regulated primarily by the Rho-ROCK-MLCKII pathway (84,88). While many molecular signatures and pathways have been investigated to understand the pathology in single cells (89,90), the mechano-mechanisms for collective migration of microtumor clusters are less understood. We utilized our previously published (18–23) 3D microtumor models, where we observed collective directional migration in large microtumors in response to the tumor-intrinsic hypoxia and radial migration in small microtumors in response to the hypoxic secretome (21). We identified key molecular signatures of collective cluster migration by analyzing RNA microarray data for these microtumors, notably differential regulation of AKT2 in hypoxia-induced directional migration and secretome-induced radial migration (23). In addition, GO term enrichment analysis of differentially expressed genes revealed upregulation of cell-cell adhesion in directional migration, while actin-microtubule cytoskeleton reorganization was upregulated in radial migration. Based on the results of the analysis, we built a multi-scale microtumor model (MSMM) focusing on the cell-cell adhesion and protrusion force to recapitulate the observed emergence of directional and radial migration.

The multi-scale microtumor model (MSMM) recapitulates both migration modes observed experimentally in microtumors and provides insight into the mechanism underlying the plasticity of tumor migration modes. MSMM provides quantitative measurements and time series for microtumor geometric features, such as tumor polarity, tumor convex area, tumor displacement distance, cell displacement distance, and classifying tumor migratory phenotypes in morphological space with machine learning methods. With the parameter scan, we found that sufficient protrusion force is necessary for tumors to migrate, consistent with the previously published reports (50,91,92). In our experiments, microtumors treated with secretome exhibited radial migration, suggesting increased protrusion force within individual cells inside the microtumors. Indeed, radial migration emerges when sufficient cellular protrusion force is generated, propelling neighboring cells to migrate collectively in diverse directions. Cellular polarity local alignment is essential for directional migration mode to emerge. Within migrating tumors, stronger cell-cell adhesion intensifies the alignment of cell polarity. This intensified alignment breaks the symmetric angular distribution of protrusion forces, leading to directional tumor migration. This result is consistent with findings on collective cluster migration that requires coordination between single cells (93,94). Our results show that radial migration emerges under sufficient protrusion force and weak adhesion, whereas directional migration emerges when local cellular polarity alignment is induced by relatively strong adhesion combined with sufficient protrusion force.

To corroborate our findings from the computational model, we conducted myosin II inhibition experiments to confirm the role of protrusion force in microtumor migration. Myosin II is a motor protein responsible for converting chemical energy into mechanical energy, generating force and movement (35,42,43,50,95,96). We observed that inhibiting myosin II reduced the radial migration of microtumors but didn’t change the migration modes, echoing our MSMM simulation results.

We then explored AKT2 as a signature regulating cell-cell adhesion and contributing to the emergence of directional migration. In microarray analysis, PIK3R3 was established as the molecular signature of directional migration in hypoxia-induced microtumors (23), a part of the PI3K-Akt pathway (68). Its downstream protein, AKT2, plays a central role in cell migration and invasion (73) and is differentially regulated in hypoxia-induced directional migration and secretome-induced radial migration (23). The pharmacological inhibition of AKT2 resulted in the transition from radial migration to directional migration mode, which was recapitulated in MSMM simulation by increasing cell-cell adhesion strength. It is technically challenging to quantify cell-cell adhesion strength directly. However, our integrated results from the ATK2 inhibition experiment and the MSMM simulations suggest that the AKT2 functions as a cell-cell adhesion regulator, thereby influencing local cell polarity alignment, which can be supported through prior literature. Indeed, AKT2 has been shown to bind to *CDH1* promoter and repress E-cadherin expression (76,77), important in maintaining cell-cell adhesion and connecting with the cortical actin cytoskeletons of cells (97,98). Strong cell-cell adhesion can contribute to directional migration due to improved cell clustering in the bloodstream (99) and more efficient cell-cell communication by passing signaling molecules to neighboring cells via gap junctions (100–102), which can help adapt gene expression to selective pressure, such as oxidative stress and fluidic shear stress in a local microenvironment (83,103,104). E-cadherin is a tumor suppressor but also functions as a promoter of growth and metastasis, based on local cellular and microenvironmental context (87,105–109). Further investigation is needed to understand the mechanistic link between AKT2 and E-cadherin to regulate cell-cell adhesion and local polarity alignment changes. Such insights can be exploited for potential therapeutic targeting.

Despite its advantages, MSMM also has disadvantages. Currently, the model does not include cell growth and cell death, but it could be implemented carefully in the future. Cell growth and death may introduce additional complexity, as Janet *et al.* mentioned that mechanical compression may correlate with invasive phenotypes (96), and mechanical compression from growth may also reduce cell growth rate (110,111). The model can be further improved by considering the effects of tumor size. Tumor size may impact the dynamics, such as intrinsic hypoxia-induced cell death in the tumor center, ROS stress signaling pathways activation, and hypoxia-driven polarity changes (112). While MSMM has the advantage of providing detailed insight into biophysical quantities that otherwise cannot be measured, such as cell-cell adhesion strength, it can be further expanded and adapted to different scenarios. For example, the model can incorporate more complex regulatory networks into its single-cell dynamics with ordinary differential equations (ODE). Overall, a thorough model would be needed to understand mechanisms in depth, such as the regulation of signal molecules and the effect of tumor size and growth rate on microtumor migration.

To summarize, we provide a mechanistic model of the interplay between protrusion force and cell-cell adhesion to understand the collective migration of microtumors using integrated experimental and computational approaches. Protrusion force is an essential driver for collective metastasis, and stronger cell-cell adhesion influences the alignment of cell polarity, leading to directional migration. An in-depth understanding of the underlying mechanisms of plasticity of tumor migration could guide the design of combination drug therapy to prevent collective migration.

## Conclusion

Collective tumor migration has been a pivotal issue in cancer research due to its complexity and many influencing factors, such as genetic regulations, microenvironmental factors, nutrient conditions, and cell-cell interactions, making its progression challenging to predict. This article presents a multi-scale microtumor model (MSMM) based on the cellular Potts model to describe and predict microtumor migration behavior.

The fundamental physical principles for the emergence of migratory modes include: 1) Sufficient protrusion force in single cells is required for microtumor migration, naturally resulting in radial migration. 2) A directional migration mode emerges in microtumors with strong cell-cell interactions and local cellular polarity alignment. The random symmetry breaking of the symmetric angular distribution of protrusion forces leads to directional tumor migration.

Experiments using microtumor models concur with our computational model’s predictions. Blebbistatin is a selective inhibitor of myosin II ATPase activity, leading to reduced protrusion force in single cells. Blebbistatin-treated microtumors show reduced radial migration, demonstrated by the decrease in the fraction of migrated tumors on day 6 and the migration distance of radially migrated tumors, consistent with MSMM. Further, we found that AKT2, the serine/threonine kinase 2, is critical for the directional migration mode and may function as a key repressor for cell-cell adhesion and local cellular polarity alignment.

MSMM and our experimental microtumor models illustrate the fundamental physical principles underlying the plasticity of tumor migration modes. This general and flexible research model allows future application in various scenarios.

## Author Contributions

SS and JX designed the experimental and computational research, respectively. HW carried out the modeling, simulations, and data analysis under the supervision of JX. CA, VA, and AJ carried out the microtumor experiments and data analysis under the supervision of SS. JVG assisted with the image analysis. YJ, WW, and JG provided advice for simulation and experiments. HW and CA wrote the first draft, and all authors edited the manuscript.

## Acknowledgment

This work is supported by the National Cancer Institute (R37 CA232209, to SS and JX), National Institute of General Medical Sciences (R01GM148525 to JX), National Science Foundation (DMS2325149 to JX), and National Eye Institute (R01EY028450) and the endowment for Frady Whipple Chair at Georgia State University (to YJ). This research is partly supported by the University of Pittsburgh Center for Research Computing through the resources provided. Specifically, this work used the H2P cluster, which is supported by NSF award number OAC-2117681. We also thank Drs. James A. Glazier, James P Sluka, T.J. Sego, and Juliano Ferrari-Gianlupi for help with Compucell3D, and Dr. GW McElfresh for help in mathematics.

