## Supplemental information for "Protrusion force and cell-cell adhesion-induced polarity alignment govern collective migration modes"

**The research was conducted at the following address:**

[1]Department of Computational & Systems Biology, School of Medicine, University of Pittsburgh, Pittsburgh, PA, USA

[2]Department of Pharmaceutical Sciences, School of Pharmacy, University of Pittsburgh, Pittsburgh, PA, USA

[3]Department of Bioengineering, Swanson School of Engineering, University of Pittsburgh, Pittsburgh, PA, USA

[4]McGowan Institute for Regenerative Medicine, University of Pittsburgh, Pittsburgh, PA, USA

[5]Department of Mathematics and Statistics, Georgia State University, Atlanta, GA, USA

[6]Department of Physics and Astronomy, University of Pittsburgh, Pittsburgh, PA, USA

[7]UPMC-Hillman Cancer Center, University of Pittsburgh, Pittsburgh, PA, USA

Present address:

[†]Department of Physics, Carnegie Mellon University, Pittsburgh, PA, USA

[#]CAS Key Laboratory for Theoretical Physics, Institute of Theoretical Physics, School of Physical Sciences, Chinese Academy of Sciences, Beijing, China

[^]School of Physical Sciences, University of Chinese Academy of Sciences, Beijing, China

[$]Department of Pharmaceutical Sciences, College of Pharmacy, University of Illinois Chicago, Chicago, IL, USA

**Local alignment of cellular polarity is crucial for the emergence of directional migration mode**

Local alignment of cellular polarity plays an important role in collective migration. In the model, the local alignment of cellular polarity is coupled with cell-cell adhesion strength. To better understand how local alignment of cellular polarity impacts the microtumor migration modes, we explored local polarity alignment as an independent variable. With force direction $\hat{f}(\sigma, t)$ defined in Eq. 13, but removing any constraints set by Eq. 14, we simulated the model for $\beta$ values of 0, 0.3, 0.6, and 1. The simulation results are shown in **Figure S4**.

Spatial velocity correlation is an essential statistical quantity in theories and models of turbulence. To understand how local polarity alignment impacts single cell dynamics in a migratory tumor, we adopt a two-point spatial velocity correlation for the final timepoint as follows (1):

$$R_{cr}(x_c, x_\sigma, t) = \frac{\langle \Delta\vec{x}_c(t) \cdot \Delta\vec{x}_\sigma(t) \rangle}{\left(\langle \Delta\vec{x}_c^2(t) \rangle \langle \Delta\vec{x}_\sigma^2(t) \rangle\right)^{\frac{1}{2}}} \qquad (S1)$$

Where $R_{cr}$ represents the spatial velocity correlation for cell pairs involving a center cell and all cells at a distance $r$; $\vec{x}_c(t)$ and $\vec{x}_\sigma(t)$ are the position vector for center cell $c$ and cells at distance $r$, as defined in Eq. 11, $\Delta\vec{x}_c(t)$ is the cell displacement vector over the past 5 MCS of center cell, and $\Delta\vec{x}_\sigma(t)$ are the cell displacement vector over the past 5 MCS of cells at a distance $r$.

Then the correlation length $d_{corr}$ is calculated by fitting the obtained $R_{cr}$ data to function:

$$R_{cr} = e^{-r/d_{corr}} \qquad (S2)$$

With $\beta = 0$, Figure S4a indicates almost no directional migration of the tumor. As $\beta$ values increase, the proportion of directionally migrating tumor cells rises significantly. High protrusion force and adhesion strength also contribute to increased spatial velocity correlation length. Indeed, local cell alignment is crucial for directional migration modes to emerge. Increasing local alignment induces directional migration mode.

**Supplementary Figures**

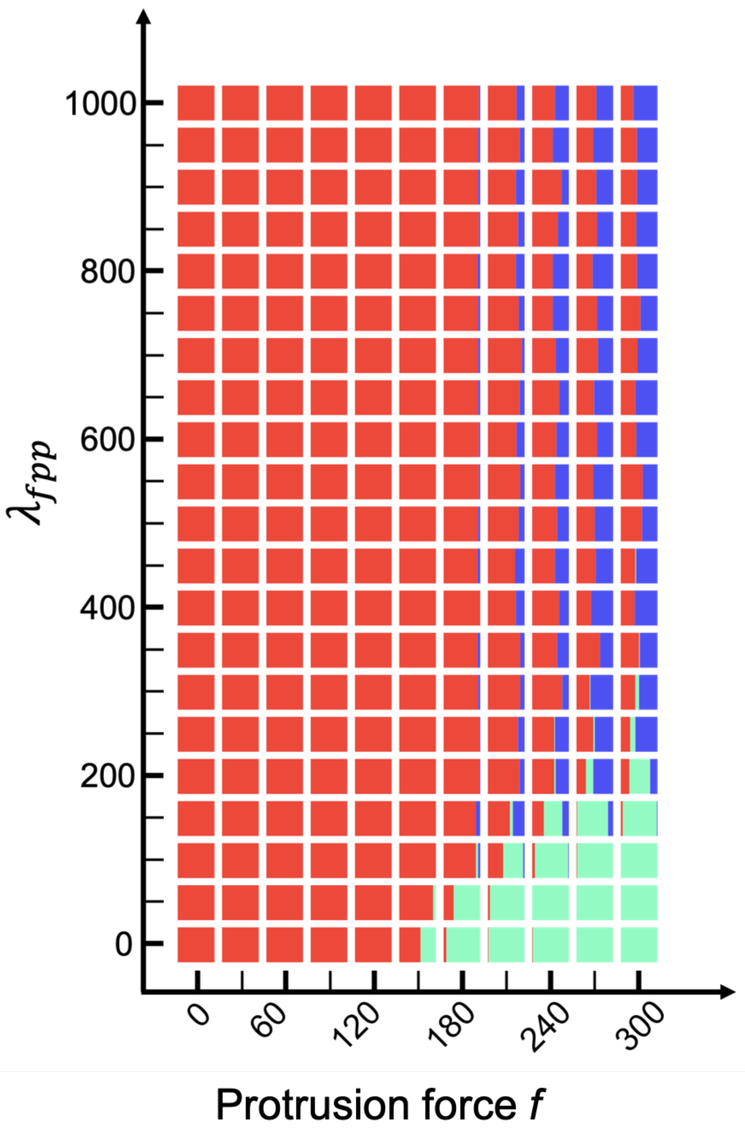

**Figure S1.** Phase graph showing tumor migration modes for parameter scan. On the x-axis, we scanned the maximum force $\eta$ for range $[0,300]$. On the y-axis, we scanned the adhesion strength $\lambda_{fpp}$ in range $[0,1000]$. The histogram is color-coded for non-migratory tumors (red), radially migrating tumors (green), and directionally migrating tumors (blue).

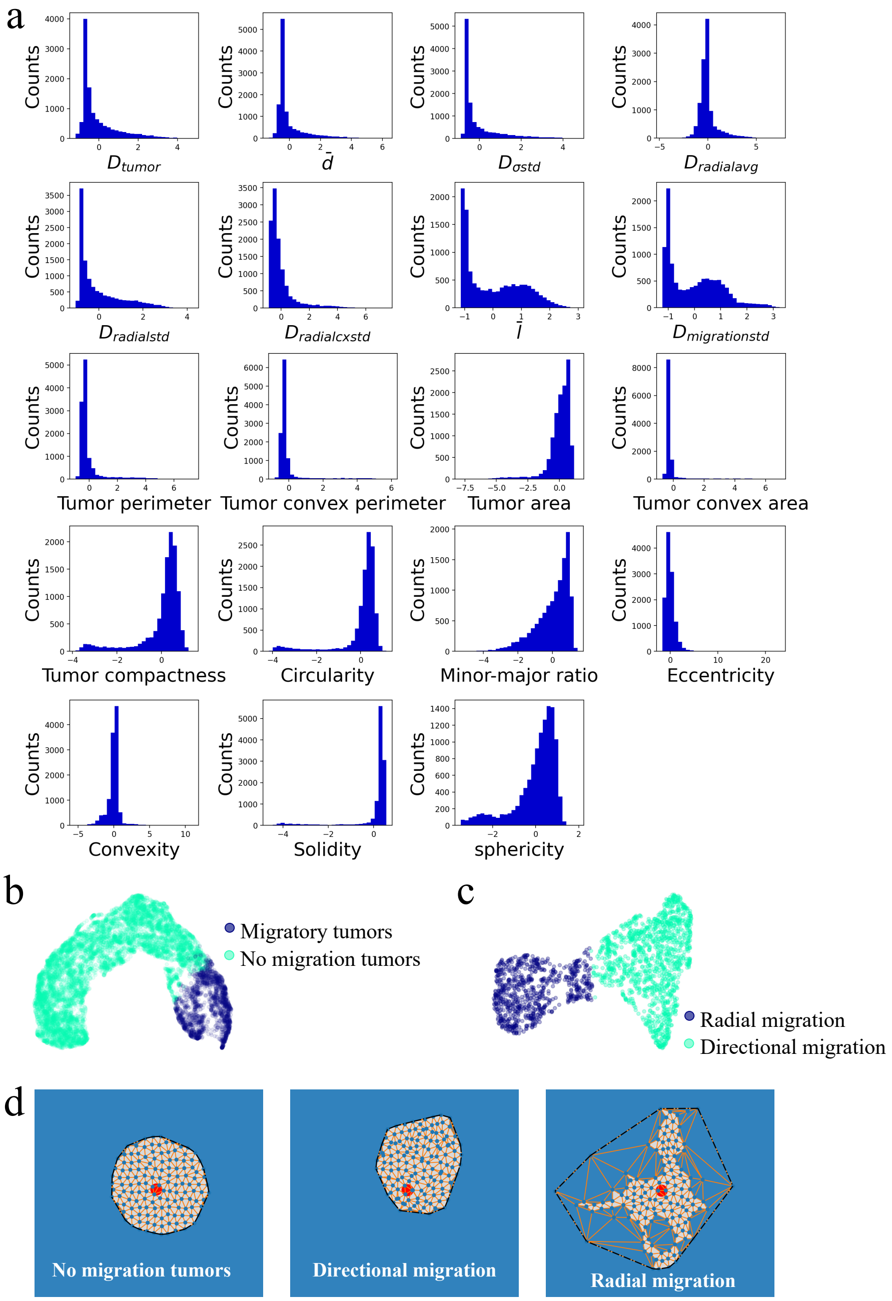

**Figure S2.** Tumor feature representations. **a.** Distribution of nineteen tumor features used for classification of tumor migration modes. **b.** UMAP representation for k-means clustering for distinguishing between non-migratory tumors and migratory tumors. **c.** UMAP representation for k-means clustering for distinguishing between radial migration and directional migration modes. **d.** Typical tumor morphologies at the final time frame for non-migratory, directionally migratory, and radially migratory tumors. Red dots are initial tumor centers. Blue dots are center coordinates for individual cells inside the tumors. Orange lines are calculated Delaunay triangles. Black dashed lines are convex hulls. The tumor area is filled with grey.

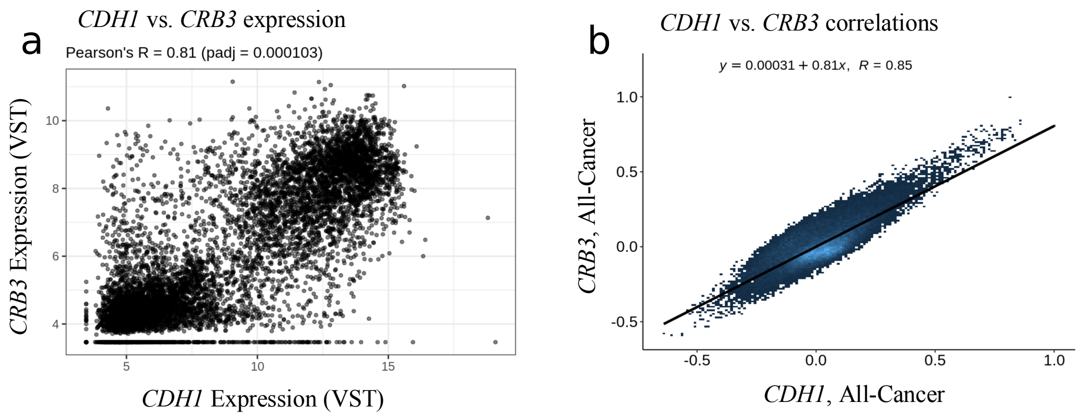

**Figure S3.** The expression of the *CDH1* gene, which encodes the key adhesion molecule E-cadherin, shows a near-linear correlation with the polarity molecule *CRB3* in cancer. **a.** Scatter plot showing gene expression (VST-transformed counts) for *CDH1* and *CRB3* in cancer tissues. **b.** Scatter plot showing the relationship between co-expression correlations for *CDH1* and *CRB3* in cancer tissues. Displayed R value determined by Pearson correlation. The figure is generated with Correlation AnalyzeR (2).

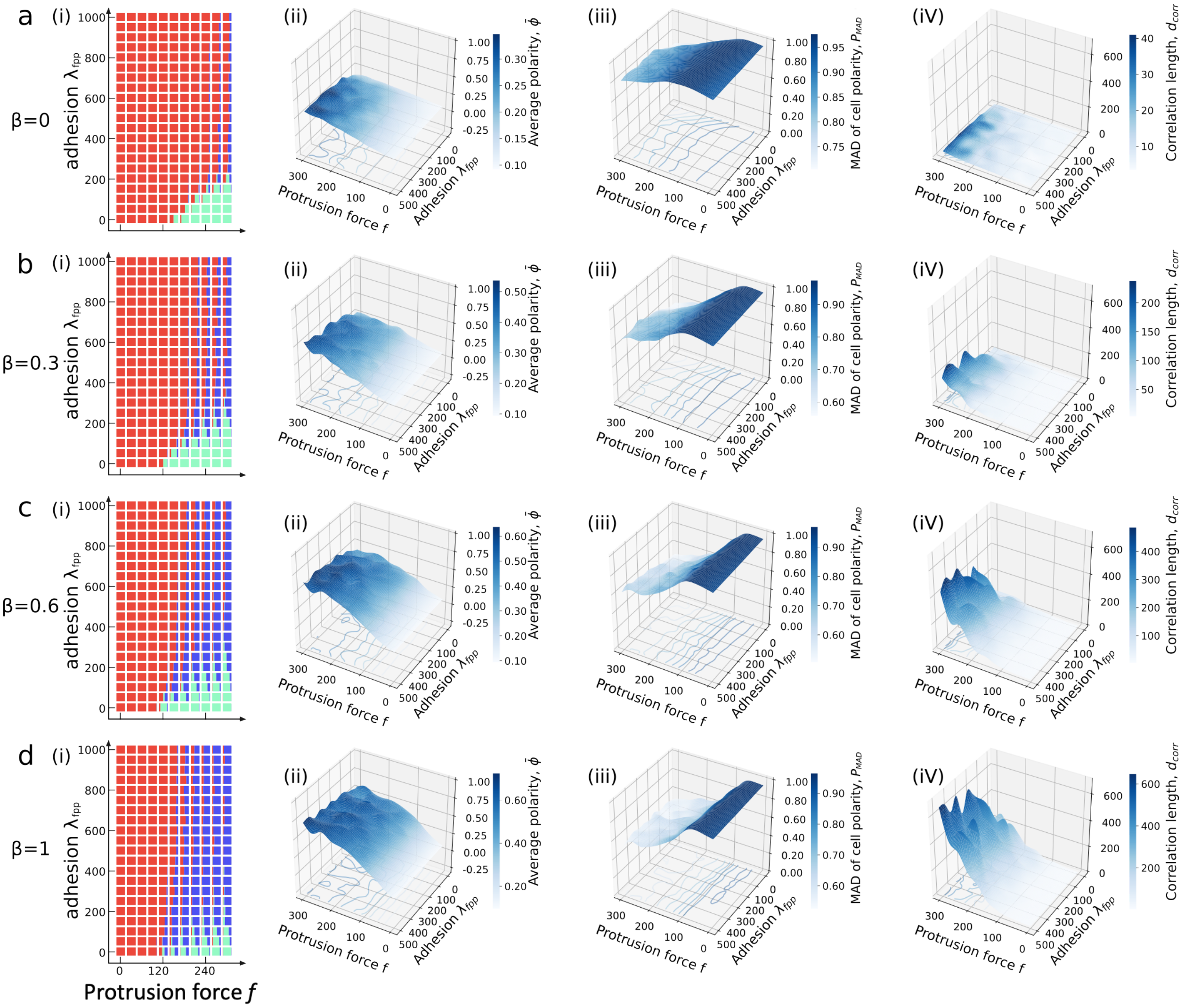

**Figure S4.** Parameter scan with varying degrees of local alignment by adjusting the coefficient $\beta$. Simulation results for $\beta$ values 0, 0.3, 0.6, and 1 are shown in panels **a**, **b**, **c**, and **d**. Microtumor migratory mode phase graphs are shown in (i). Microtumor properties, such as tumor average polarity $\bar{\phi}$, tumor average cell polarity $P_{MAD}$ and correlation length at the end timepoint $d_{corr}$ are shown in (ii), (iii) and (iv).

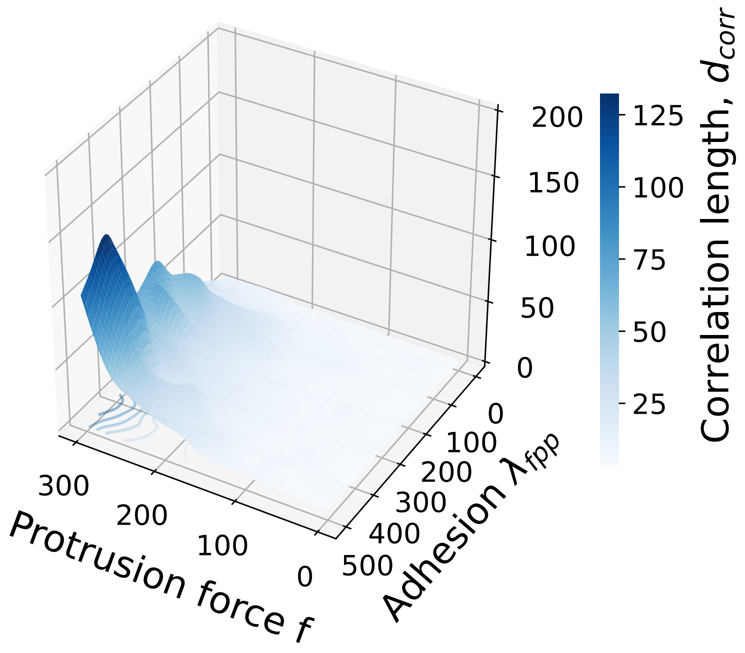

**Figure S5**. The relationship between protrusion force and cell-cell adhesion on spatial velocity correlation length. The plot illustrates how varying levels of protrusion force ($f$) and cell-cell adhesion ($\lambda_{fpp}$) influence the correlation length ($d_{corr}$). As both protrusion force and adhesion increase, the spatial velocity correlation length also increases, indicating enhanced coordination in cell movement.

**Supplementary Videos**

**Video S1.** Movie of the *in silico* MSMM model recapitulating radial migration with $f = 150$ and $\lambda_{fpp} = 20$, corresponding to the data presented in Figure 3a (i). Real time effective force $\hat{F}$ is color coded.

**Video S2.** Movie of the *in silico* MSMM model recapitulating directional migration with $f = 300$ and $\lambda_{fpp} = 400$ corresponding to the data presented in Figure 3a (ii). Real time effective force $\hat{F}$ is color coded.

### Supplementary Tables

#### Table S1. Multi-scale microtumor model parameters.

| Multi-scale microtumor model parameters | | |
|---|---|---|
| **Parameter** | **Definition** | **Values** |
| Dimensions | Lattice dimensions | 256×256 |
| $T$ | Temperature | 25 |
| $J_{medium,medium}$ | Bond energy (medium-medium) | -1 |
| $J_{medium,cell}$ | Bond energy (medium-cell) | -2 |
| $J_{cell,cell}$ | Bond energy (cell-cell) | -6 |
| $\lambda_{area}$ | Cell area constraint | 4 |
| $A_{\sigma}$ | Cell target area | 50.24 |
| $\lambda_{peri}$ | Cell perimeter constraint | 4 |
| $L_{\sigma}$ | Cell target perimeter | 25 |
| $\lambda_{fpp}$ | Coefficient regulating cell-cell adhesion strength | [0, 1000] |
| $D_{\sigma\omega}$ | Neutral cell pair distance where cell-cell adhesion energy is 0 | 8 pixel |
| $s_{medium}$ | Constant secretome field in the medium | 1/pixel |
| $D$ | Global secretome field diffusion constant | 1 |
| $k_{uptake}$ | Cell (and protrusion cell) secretome relative uptake | 0.0025/pixel |
| $\zeta$ | Coefficient for converting secretome uptake signal to production rate of R | Assigned during simulation |
| $\beta_{max}$ | Neighbor sharing polarity constraint | 0.4 |

**Table S2. Parameter table of ODE dynamics for protrusion force in single cells.**

| Parameter table of ODE dynamics for protrusion force in single cells | | | |
|---|---|---|---|
| **Parameter** | **Definition** | **Values** | **Reference** |
| $b$ | Basal production rate protrusion force regulator, R | $1\ \mu M/s$ | |
| $s$ | Production rate of secretome-induced R. This parameter has the highest value at the tumor boundary. | $4\ \mu M/s$ | |
| $d$ | Degradation rate of protrusion force regulator, R | $0.1/s$ | Estimated within reasonable range (3-6) |
| $f$ | Maximum protrusion force | [0, 300] | |
| $K_R$ | Half saturation concentration | $40\ \mu M$ | Estimated within reasonable range (7) |

**Table S3. Geometric features were used for clustering the tumor migration modes.**

<table>
<tr><th colspan="6">Geometric features were used for clustering the tumor migration modes</th></tr>
<tr><th>#</th><th>Geometric features</th><th>Symbol</th><th>Formula</th><th>Annotation</th><th>Type</th></tr>
<tr><td>1</td><td>Displacement distance for the center of tumor mass</td><td>$D_{tumor}$</td><td>$D_{tumor} = \left| \frac{\sum_{\sigma=1}^{N} \vec{x}_\sigma}{N} - \vec{\varepsilon} \right|$</td><td rowspan="8">$\vec{x}_\sigma$ is the cell $\sigma$ position vector at the final time frame (270 MCS*).<br><br>$\vec{x}_{\sigma init}$ is the cell $\sigma$ position vector at the initial time frame (90 MCS).<br><br>$\vec{\varepsilon}$ represent a vector for the initial tumor center, at pixel position $(128,128)$.<br><br>$N$ is the number of cells in the tumor.<br><br>$M$ is the number of cells on the boundary of the tumor.<br><br>$d_\sigma$ is the distance for cell $\sigma$ position to the initial tumor center.<br><br>$l_\sigma$ is the distance for cell $\sigma$ displacement compared to its initial position.</td><td rowspan="8">Distance measures</td></tr>
<tr><td>2</td><td>Average distance of displacement for cells</td><td>$\bar{d}$</td><td>$d_\sigma = |\vec{x}_\sigma - \vec{\varepsilon}|$<br>$D_{\sigma avg} = \frac{\sum_{\sigma=1}^{N} d_\sigma}{N}$</td></tr>
<tr><td>3</td><td>Standard deviation of $d_\sigma$</td><td>$D_{\sigma std}$</td><td>$D_{\sigma std} = \sqrt{\frac{\sum_{\sigma=1}^{N}(d_\sigma - \bar{d})^2}{N}}$</td></tr>
<tr><td>4</td><td>Average radial distance measure</td><td>$D_{radialavg}$</td><td>$D_{radialavg} = \frac{\sum_{\sigma=1}^{M} d_\sigma}{M}$</td></tr>
<tr><td>5</td><td>Standard deviation of radial distance measure</td><td>$D_{radialstd}$</td><td>$D_{radialstd} = \sqrt{\frac{\sum_{\sigma=1}^{M}(d_\sigma - \bar{d})^2}{M})}$</td></tr>
<tr><td>6</td><td>Standard deviation of radial distance measure comparing to real time tumor center</td><td>$D_{radialcxstd}$</td><td>$r_\sigma = \vec{x}_\sigma - \frac{\sum_{\sigma=1}^{N} \vec{x}_\sigma}{N}$<br>$D_{radialcxstd} = \sqrt{\frac{\sum_{\sigma=1}^{M}(r_\sigma - \bar{r})^2}{M}}$</td></tr>
<tr><td>7</td><td>Average cell migratory distance compared to the initial state</td><td>$\bar{l}$</td><td>$l_\sigma = \vec{x}_\sigma - \vec{x}_{\sigma init}$<br>$\bar{l} = \frac{\sum_{\sigma=1}^{N} l_\sigma}{N}$</td></tr>
<tr><td>8</td><td>Standard deviation of cell migratory distance compared to initial position.</td><td>$D_{migrationstd}$</td><td>$D_{migrationstd} = \sqrt{\frac{\sum_{\sigma=1}^{N}(l_\sigma - \bar{l})^2}{N}}$</td></tr>
<tr><td>9</td><td>Tumor perimeter</td><td>$P_{tumor}$</td><td>The number of pixels on the edges of the tumor boundary, defined by Delaunay triangles.</td><td></td><td>Shape measures</td></tr>
</table>

| | | | | |
|---|---|---|---|---|
| 10 | Tumor convex perimeter | $P_{convex}$ | The number of pixels on the edges of the convex hull. | |
| 11 | Tumor area | $A_{tumor}$ | $A_{tumor} = \frac{\# \, of \, pixels \, in \, tumor}{256 \times 256}$ | For simplicity, the tumor area is normalized by the simulation panel size |
| 12 | Tumor convex area | $A_{convex}$ | $A_{convex} = \frac{\# \, of \, pixels \, in \, convex \, hull}{256 \times 256}$ | For simplicity, the tumor convex area is normalized by the simulation panel size |
| 13 | Tumor compactness | $F_{compact}$ | $F_{compact} = \frac{4\pi \cdot A_{tumor}}{P_{tumor}^2}$ | |
| 14 | Tumor circularity | $F_{circularity}$ | $F_{circular} = \frac{4\pi \cdot A_{tumor}}{P_{convex}^2}$ | $P_{convex}$ is the tumor perimeter calculated by the convex hull algorithm. |
| 15 | Minor-major ratio | $F_{minormajor}$ | $F_{minormajor} = \frac{D_{minor}}{D_{major}}$ | $D_{minor}$ is the length of the minor axis of the ellipse that has the same normalized second central moments as the region. Similarly, $D_{major}$ is the length for the major axis. |
| 16 | Eccentricity | $F_{eccentricity}$ | For an ellipse that has the same second moments as the tumor region, eccentricity is the ratio of the focal distance over the major axis length $D_{major}$. | |
| 17 | Convexity | $F_{convexity}$ | $F_{convexity} = \frac{P_{convex}}{P_{tumor}}$ | |
| 18 | Solidity | $F_{solidity}$ | $F_{solidity} = \frac{A_{tumor}}{A_{convex}}$ | |
| 19 | Sphericity | $F_{sphericity}$ | $F_{sphericity} = \frac{R_{inscribing}}{R_{circumscribing}}$ | $R_{inscribing}$ and $R_{circumscribing}$ are the radius of circles inscribing and circumscribing the tumor, respectively. |

Note: MCS*: Monte Carlo step
